# Pathway enrichment analysis of -*omics* data

**DOI:** 10.1101/232835

**Authors:** Jüri Reimand, Ruth Isserlin, Veronique Voisin, Mike Kucera, Christian Tannus-Lopes, Asha Rostamianfar, Lina Wadi, Mona Meyer, Jeff Wong, Changjiang Xu, Daniele Merico, Gary D. Bader

**Affiliations:** Computational Biology Program, Ontario Institute for Cancer Research, Toronto, Ontario, Canada; Department of Medical Biophysics, University of Toronto, Toronto, Ontario, Canada; The Donnelly Centre, University of Toronto, Toronto, Ontario, Canada; Deep Genomics Inc., Toronto, Ontario, Canada; The Centre for Applied Genomics (TCAG), The Hospital for Sick Children, Toronto, Ontario, Canada; Department of Molecular Genetics, University of Toronto, Toronto, Ontario, Canada; Department of Computer Science, University of Toronto, Toronto, Ontario, Canada

**Keywords:** bioinformatics, genomics, pathway enrichment analysis, functional analysis, network analysis, Cytoscape, Enrichment Map, g:Profiler, GSEA

## Abstract

Pathway enrichment analysis helps gain mechanistic insight into large gene lists typically resulting from genome scale (–*omics*) experiments. It identifies biological pathways that are enriched in the gene list more than expected by chance. We explain pathway enrichment analysis and present a practical step-by-step guide to help interpret gene lists resulting from RNA-seq and genome sequencing experiments. The protocol comprises three major steps: define a gene list from genome scale data, determine statistically enriched pathways, and visualize and interpret the results. We focus on differentially expressed genes and mutated cancer genes, however the described principles can be applied to diverse –*omics* data. The protocol is designed for biologists with no prior bioinformatics training and uses freely available software including g:Profiler, GSEA, Cytoscape and Enrichment Map.

## INTRODUCTION

Comprehensive surveys of DNA, RNA and proteins in biological samples are now routine. The resulting data are growing exponentially and their analysis helps discover novel biological functions, genotype-phenotype relationships and disease mechanisms. However, analysis and interpretation of these data is a major challenge for many researchers. Analyses often result in long lists of genes that require an impractically large amount of manual literature searching to interpret. A standard approach to addressing this problem is pathway enrichment analysis, which summarizes the large gene list as a smaller list of more easily interpretable pathways. Pathways are statistically tested for over-representation in the experimental gene list above what is expected by chance. For instance, experimental data containing 40% cell cycle genes is surprisingly enriched given that only 8% of human protein-coding genes are involved in this process.

In a recent example, we used pathway enrichment analysis to help identify histone and DNA methylation by the Polycomb repressive complex (PRC2) as the first rational therapeutic target for ependymoma, one of the most prevalent childhood brain cancers^1^. This pathway is targetable by available drugs, such as 5-azacytidine, which was used on a compassionate basis in a terminally ill patient and stopped rapid metastatic tumour growth. In another example, we analysed rare copy number variants (CNVs) in autism and identified several significant pathways affected by gene deletions, whereas only few significant hits were identified with case-control association tests of single genes or loci^2,3^. These examples illustrate the useful insights into biological mechanisms that can be achieved using pathway enrichment analysis.

This protocol covers pathway enrichment analysis of large gene lists typically derived from genome scale (“–*omics*”) technology. The protocol is intended for experimental biologists who are interested in interpreting their –*omics* data. It requires only an ability to learn and use “point-and-click” computer software, although advanced users can benefit from automatic analysis scripts we provide. We analyse human gene expression and somatic mutation data as examples, however our conceptual framework is applicable to analysis of lists of genes or biomolecules from any organism derived from large-scale data, including proteomics, genomics, epigenomics and gene regulation studies. The protocol uses free, easy to use, updated and well documented software (g:Profiler^4^, GSEA^5^, Cytoscape^6^, Enrichment Map^7^). We next present the major steps of pathway enrichment analysis, describe each in detail and then provide a detailed step-by-step protocol.

## Protocol overview

Pathway enrichment analysis involves three major steps (**Figure 1; See Box 1 for basic definitions).**

1. **Define a gene list of interest using** –*omics* **data.** An –*omics* experiment comprehensively measures the activity of genes in an experimental context. The raw data generally require computational processing, such as normalization and scoring to identify genes of interest, considering the experimental design. For example, a list of genes differentially expressed between two groups of samples can be derived from RNA-seq data.
2. **Perform pathway enrichment analysis.** A statistical method is used to identify pathways enriched in the gene list from step 1, relative to what is expected by chance. All pathways in a given database are tested for enrichment in the gene list. Several established pathway enrichment analysis methods are available and the choice of which to use depends on the type of gene list.
3. **Visualize and interpret pathway enrichment analysis results.** Many enriched pathways may be identified in step 2, often including related versions of the same pathway. Visualization can help identify the main biological themes and their relationships in this list for in-depth study.

**Figure 1.**
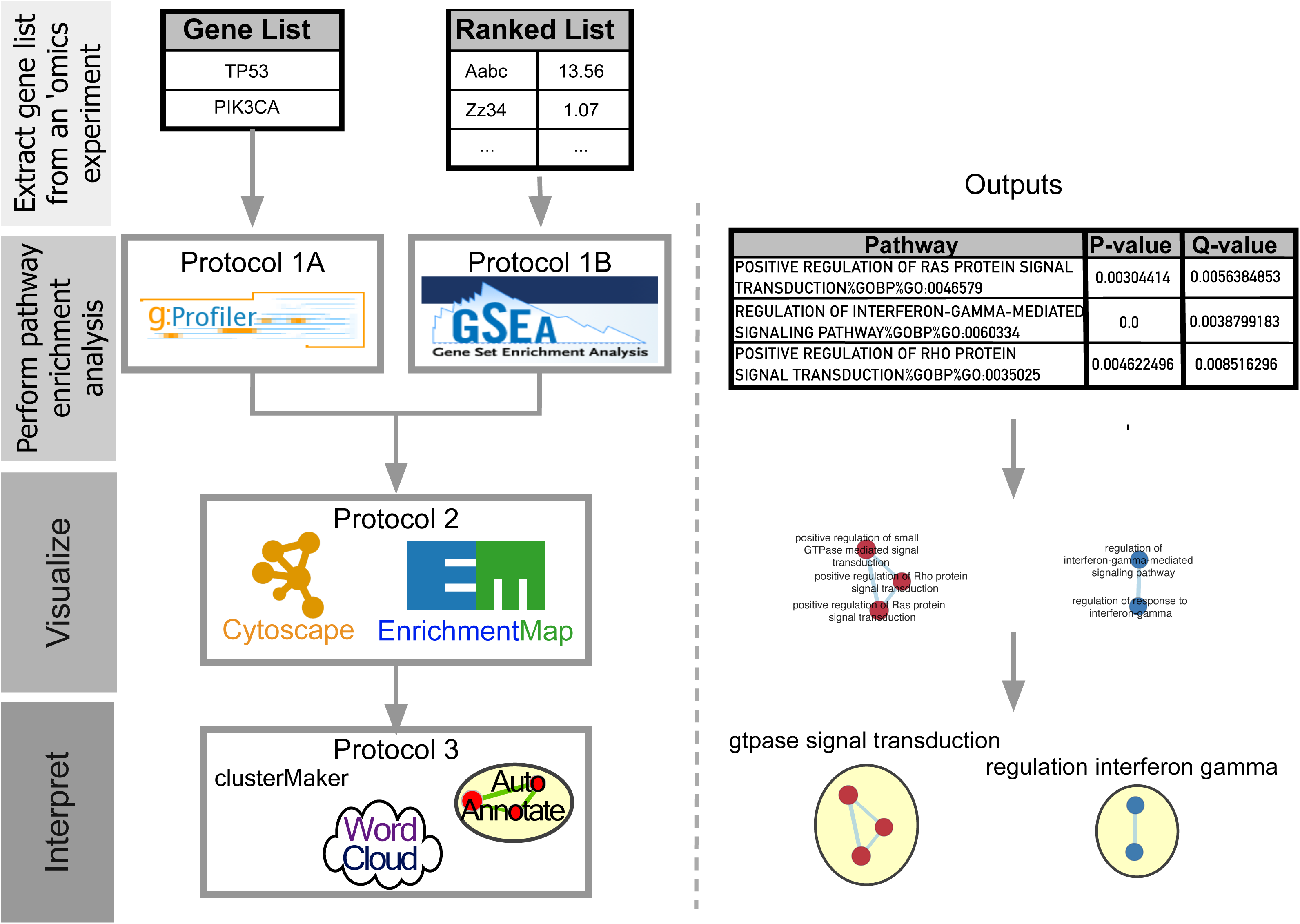
Protocol overview. Gene lists derived from diverse ‘omics data undergo pathway enrichment analysis, using g:Profiler or GSEA, to identify pathways that are enriched in the experiment. Pathway enrichment analysis results are visualized to aid interpretation (using Cytoscape and associated apps Enrichment Map, AutoAnnotate, Word Cloud and clusterMaker2). Protocol overview is shown on the left, starting from gene list input, and example outputs at each stage are shown on the right.

### Step 1: Define a gene list of interest using -omics data

Genome-scale experiments generate raw data that must be processed to obtain gene-level information suitable for pathway enrichment analysis. The specific processing steps are particular to the –*omics* experiment type and may be standard or complex. Standard processing methods are available for established –*omics* technologies that are most conveniently performed by the core facility that generates the data. Standard protocols are available, for example for RNA-seq^8^, microarrays^9^, protein expression^10^, genomic variant annotation^11^, and DNA methylation^12^.

There are two major ways to define a gene list from –*omics* data: list or ranked list. Certain –*omics* data naturally produce a gene list, such as all somatically mutated genes in a tumor from exome sequencing, or all proteins that interact with a bait in a proteomics experiment. Such a list is suitable for direct input into pathway enrichment analysis using g:Profiler protocol 1A. Other –*omics* data naturally produce ranked lists. For example, a list of genes can be ranked by differential gene expression score or sensitivity in a genome wide CRISPR screen. Early pathway enrichment analysis approaches involved applying a threshold to a ranked gene list (e.g. FDR-adjusted p-value below 0.05 and fold-change above 2); however, this is often arbitrary and thus not recommended, especially when meaningful ranks are available for all or most of the genes in the genome. Modern approaches, like GSEA, are designed to analyze ranked lists of all available genes and do not require a threshold. A ranked list is suitable for input into pathway enrichment analysis using GSEA protocol 1B. Alternatively, a partial ranked gene list can be analysed using g:Profiler.

As an example, we describe analysis of raw RNA-seq data to define a ranked gene list. DNA sequence reads are quality filtered (e.g. by trimming to remove low quality bases) and mapped to a genome-wide reference set of transcripts to enable counting reads per transcript. Read counts are aggregated at the gene level (counts per gene). Typically, RNA-seq data for multiple biological replicates (three or more) for each of multiple experimental conditions (two or more, e.g. treatment vs. control) are available (**Box 2 – experimental design**). Read counts per gene are normalized across all samples to remove unwanted technical variation between samples, for example, due to differences in sequencing lane or total read number per sequencing run^13-15^. Next, read counts per gene are tested for differential expression across sample groups (e.g. treatment vs. control) (**Supplementary Protocol 1**). Software packages such as edgeR^16^, DESeq^17^, limma^18^, and Cufflinks^19^ implement procedures for RNA-seq data normalization and differential expression analysis. Differential gene expression analysis results include: 1) the p-value of the significance of differential expression; 2) the related q-value (a.k.a adjusted p-value) that has been corrected for multiple testing across all genes (e.g. using the Benjamini-Hochberg False Discovery Rate (FDR) procedure^20^); 3) effect size and direction of expression change (expressed as fold-change or log-transformed fold-change) so that up-regulated genes are positive and at the top of the list and down-regulated genes are negative and at the bottom of the list. The list of all genes is then ranked by one or more of these values (e.g. −log10 p-value multiplied by the sign of log-transformed fold-change). This ranked list is provided as input to pathway enrichment analysis (ranked gene list, no threshold needed, **pathway enrichment analysis using GSEA protocol 1B**).

### Step 2A: Pathway enrichment analysis of a gene list using g:Profiler

The default analysis implemented in g:Profiler and similar web-based tools *(e.g.,* Panther^21^, ToppGene^22^, Enrichr^23^, DAVID^24^) searches for pathways whose genes are significantly enriched (i.e. over-represented) in the fixed list of genes of interest, compared to all genes in the genome. The p-value of the enrichment is computed using a Fisher’s exact test and multiple test correction is applied (**Box 3**).

The g:Profiler tool also includes an ordered enrichment test, which is suitable for lists of up to a few thousand genes that are ordered by a score, while the rest of the genes in the genome lack meaningful signal for ranking. For example, significantly mutated genes may be ranked by a score from a cancer driver prediction method^25^. This analysis repeats a modified Fisher’s exact test on incrementally larger sub-lists of the input genes and reports the sub-list with the strongest enrichment p-value for every pathway^26^. g:Profiler searches a set of pathway, network, regulatory motif, and phenotype gene sets. Major gene set categories can be selected to customize the search.

Pathway enrichment methods that use the Fisher’s exact test, or related overrepresentation tests, require the definition of a set of background genes for comparison. All annotated protein-coding genes are often used as default. This leads to inappropriate inflation of p-values and false positive results if the experiment can directly measure only a subset of all genes. For example, setting a custom background is important in analysing data from targeted sequencing or phosphoproteomics experiments. The appropriate custom background would include all genes in the sequencing panel or all known phosphoproteins, respectively.

### Step 2B: Pathway enrichment analysis of a ranked gene list using GSEA

Pathway enrichment analysis of a ranked gene list is implemented in the GSEA algorithm^5^. GSEA is a threshold-free method that analyzes all genes based on their differential expression rank, or other score, without prior gene filtering. GSEA is particularly suitable and recommended when ranks are available for all or most of the genes in the genome (e.g. for RNA-seq data), however it is limited or inapplicable when only a small portion of genes have ranks available.

The GSEA method searches for pathways whose genes are enriched at the top or bottom of the ranked gene list, more so than expected by chance alone. For instance, if the top most differentially expressed genes are involved in the cell cycle, this suggests that the cell cycle pathway is regulated in the experiment. In contrast, the cell cycle pathway is likely not significantly regulated if the cell cycle genes appear randomly scattered through the whole ranked list. To calculate an enrichment score (ES) for a pathway, GSEA progressively examines genes from the top to the bottom of the ranked list, increasing the enrichment score if a gene is part of the pathway and decreasing the score otherwise. These running sum values are weighted, so that enrichment in the very top-(and bottom-) ranking genes is amplified, whereas enrichment in genes with more moderate ranks are not amplified. The ES score is calculated as the maximum value of the running sum and normalized relative to pathway size, resulting in a normalized enrichment score (NES) that reflects the enrichment of the pathway in the list. Positive and negative NES values represent enrichment at the top and bottom of the list, respectively. Finally, a permutation-based p-value is computed and corrected for multiple testing to produce a permutation-based FDR *q*-value that ranges from zero (highly significant) to one (not significant) (**Box 3**). The same analysis is performed starting from the bottom of the ranked gene list to identify pathways enriched in the bottom of the list. Resulting pathways are selected using the FDR *q*-value threshold (e.g. *q*<0.05), and ranked using NES. It is also useful to inspect the “leading edge” genes that contribute to the increase of the enrichment score before it peaks.

GSEA has two methods to determine the statistical significance of the enrichment score and compute a p-value: gene set permutation and phenotype permutation. For gene set permutation, input is a ranked list and GSEA compares the observed pathway enrichment score to a distribution of scores obtained by repeating the analysis with randomly sampled gene sets of matching sizes (e.g. 1,000 times). In the phenotype permutation mode, input is expression data for all samples along with a definition of sample groups (called ‘phenotypes’ - e.g. cases vs. controls, tumor vs. normal) to be compared against each other. The observed pathway enrichment score is compared to a distribution of scores obtained by randomly shuffling the samples among phenotype categories and repeating the analysis (e.g. 1,000 times), including computation of the ranked gene list and resulting pathway enrichment score. The gene set permutation mode is recommended for studies with limited variability and biological replicates (i.e. 2 to 5 per condition). In this case, differential gene expression analysis should be computed using methods that include variance stabilization, outside of GSEA. If more replicates are available (above 6 to 10 per condition), the phenotype permutation should be used, offering as a main advantage that it models gene correlations in the gene expression matrix, unlike the gene set permutation approach. This protocol only covers gene set permutation because it can be accomplished using easy to use GSEA software, whereas phenotype permutation for RNA-seq data requires computing the enrichment score and differential expression statistics on thousands of phenotype randomizations, which currently requires custom programming outside of GSEA.

By default, the GSEA desktop software searches the MSigDB gene set database that includes pathways, published gene signatures, microRNA target genes and other gene set types (**Box 4**). The user can also provide a custom database as a text-based ‘Gene Matrix Transposed’ (GMT) file where each line defines a pathway, with its name, identifier and a list of gene identifiers that match the input gene list.

### General recommendations for pathway enrichment analysis

We recommend searching enrichment only of pathway gene sets at first, as these capture familiar normal cellular processes that are easy to interpret. Gene Ontology (GO)^27^ biological process terms and manually curated molecular pathways from Reactome^28^, Panther^21^, HumanCyc ^29^, and NetPath^30^ are good resources for human pathways (**Box 4**). GO biological process annotations include a mix of manually curated and electronically inferred sources. We recommend excluding those with the lower quality ‘inferred from electronic annotation’ (IEA) evidence code, unless no enriched pathways are found. Pathway definitions change rapidly and it is essential to use updated databases of gene annotations as outdated databases can lead to missed discoveries^31^.

Different types of gene sets help answer a variety of questions. For instance, gene sets corresponding to microRNA and transcription factor targets can be used to discover important regulators^32,33^. The use of additional sets must be carefully considered, as simultaneously analyzing all available gene sets increases the number of statistical tests and leads to more conservative p-values following multiple test correction (**Box 3**).

Gene set size is important to consider. Small pathways (e.g. less than ten or fifteen genes) should be excluded because these are often numerous, negatively affecting multiple test correction, and redundant with larger pathways. For human gene expression analysis, large pathways (e.g. over 300 genes) should also be excluded as these are overly general (e.g. ‘metabolism’) and don’t contribute to interpretability of results. However, for other gene set types and organisms, larger pathways and gene sets may need to be included (e.g. up to 1000 genes).

A pathway enrichment analysis resulting in few or no enriched pathways may be caused by suboptimal statistical processing used to define the gene list. If the gene list ranks are too noisy (interfering with the signal of having the most important genes at the top of the list), all or no genes are highly significant, then enriched pathways are unlikely to be found. If the gene list has been correctly defined, increasing the number of pathways and gene sets searched or setting more liberal filters may improve results. Finally, pathway enrichment analysis results can change based on the parameters used (e.g. minimum and maximum pathway size or selected pathway databases), thus the robustness of conclusions should be tested by varying these parameters.

### Step 3: Visualising and interpreting pathway enrichment analysis results

Pathway information is inherently redundant, as genes often participate in multiple pathways, and some pathway databases organize pathways hierarchically by including general and specific pathways with many shared genes (e.g. ‘cell cycle’ and ‘M-phase of cell cycle’). Pathway enrichment analysis often highlights several versions of the same pathway as a result. Collapsing redundant pathways into a single biological theme simplifies interpretation. We recommend addressing such redundancy with the Enrichment Map visualization method^7^ or similar^34^. An enrichment map is a network representing overlaps among enriched pathways. Pathways are represented as circles (nodes) that are colored by enrichment score and are connected with lines (edges) sized based on the number of genes shared by the connected pathways. Network layout and clustering algorithms are used to automatically display and group similar pathways as major biological themes (**Figure 1**). The Enrichment Map software takes as input a text file containing pathway enrichment analysis results and another text file containing the pathway gene sets used in the original enrichment analysis. Interactive exploration of pathway enrichment score (filtering nodes) and connections between pathways (filtering edges) is possible (see **visualize enrichment results with Enrichment Map, protocol 2**). Multiple enrichment analysis results can be simultaneously visualized in a single enrichment map, in which case different colors are used on the nodes for each enrichment. If the gene expression data are optionally loaded, clicking on a pathway node will display a gene expression heat map of all genes in the pathway.

An enrichment map helps identify interesting pathways and themes. First, expected themes should be identified to help validate the pathway enrichment analysis results (positive controls). For instance, growth related pathways are expected to be identified in cancer samples relative to controls. Second, pathways not previously associated with the experimental context are evaluated more carefully as potential discoveries. Pathways and themes with the strongest enrichment scores should be studied first, followed by progressively weaker signals (see **navigating and interpreting the Enrichment Map, protocol 3**). Third, interesting pathways are examined in more detail, examining genes within the pathways (e.g. expression heat maps and the GSEA leading edge genes). Further, gene expression values can be overlaid on a pathway diagram, if available, from databases such as Pathway Commons^35^, Reactome^28^, KEGG^36^ or WikiPathways^37^ using tools such as PathVisio^38^. If a diagram is not available, tools such as STRING^39^ or GeneMANIA^40^ can be used with Cytoscape^6^ to define an interaction network among pathway genes for expression overlay. This helps visually identify pathway components (e.g. branches or single elements) that are most altered (e.g., differentially expressed) in the experiment. Additionally, master regulators for enriched pathways can be searched for by integrating miRNA^32^ or transcription factor^33^ target gene sets using the Enrichment Map post-analysis tool. Finally, pathway enrichment analysis results can be published to support a scientific conclusion (e.g. functional differences of two cancer subtypes), used for hypothesis generation or planning experiments to support the identification of novel pathways.

### Caveats of pathway enrichment analysis

The following caveats are important to consider when interpreting pathway enrichment analysis results.

- Pathway enrichment analysis assumes that a strong experimental signal of pathways reflects the biology addressed by the experiment. For instance, in a transcriptomics experiment, we assume that evolution has optimized a cell to express a pathway only when needed and these can be identified. Pathway activity not controlled by gene expression (e.g. post-translational regulation) will not be observed.
- Unexpected biological themes may indicate problems with experimental design, data generation or analysis. For example, enrichment of the apoptosis pathway may indicate a problem with the experimental protocol that led to increased cell death during sample preparation. In these cases, the experimental design and data generation should be carefully reviewed prior to pathway analysis.
- Pathway databases, and therefore enrichment results are biased towards well known pathways.
- Multi-functional genes that are highly ranked in the gene list may lead to enrichment of many different pathways, some of which are not relevant to the experiment. Repeating the analysis after excluding such genes may reveal pathways whose enrichment is overly-dependent on their presence or confirm the robustness of pathway enrichment.
- Pathway enrichment analysis ignores genes with no pathway annotations, sometimes called “dark matter of the genome”, and these genes should be studied separately.
- Most enrichment analysis methods make unrealistic assumptions of statistical independence among genes as well as pathways. Some genes may be always co-expressed (e.g. genes within a protein complex) and some pathways have genes in common. Thus, standard false discovery rates, which assume statistical independence between tests, are often either more or less conservative than ideal. Nonetheless, they should still be used to adjust for multiple testing and rank enriched pathways for exploratory analysis and hypothesis generation. Custom permutation tests may lead to better estimates of false discovery (**Box 3**).
- By representing pathways as gene sets, many biological details such as proteinprotein interactions, biochemical reactions, post-translational modifications, protein complexes, and activation and inhibition relationships are ignored. These issues are addressed by advanced methods that consider mechanistic pathway details, however this is still an active area of research (**Box 5**).

### Working with diverse –*omics* data

Pathway enrichment analysis is generally applicable to any experiment that can generate a list of genes, though experiment specific issues must be considered:

- Genes are associated with many, diverse database identifiers (IDs). We recommend using unambiguous, unique and stable IDs, as some IDs become obsolete over time. For human genes, we recommend using the Entrez Gene database IDs (e.g. *4193* corresponds to *MDM2,* http://www.ncbi.nlm.nih.gov/gene/4193) or gene symbols *(MDM2* is the official symbol recommended by the HUGO Gene Nomenclature Committee). As gene symbols change over time, we recommend maintaining both gene symbols and Entrez Gene IDs. We recommend UniProt accession numbers for proteins (e.g. *Q00987* for MDM2, http://www.uniprot.org/uniprot/Q00987) and Human Metabolome Database (HMDB) IDs for metabolites (e.g. ATP is denoted as *HMDB00538,* http://www.hmdb.ca/metabolites/HMDB00538). The g:Profiler and related g:Convert tool support automatic conversion of multiple ID types to standard IDs.
- Pathway enrichment analysis of short non-coding genomic regions such as transcription factor binding sites from ChIP-seq experiments need additional consideration. Genomic regions must be mapped to protein-coding genes and corrected for biases such as increased signal in longer genes. Tools such as GREAT^41^ automatically perform both tasks.
- Large genomic intervals that span multiple genes (e.g. from genome-wide associations, copy number variation and differentially methylated regions) require specialized tests such as the PLINK CNV gene set burden test^42^ or INRICH^43^. Standard enrichment tests often reveal genes clustered in the genome that are strongly statistically inflated due to incorrectly counting each gene as an independent signal. These include olfactory receptors, histones, major histocompatibility complex (MHC) members and homeobox transcription factors. A simple solution involves selecting only one representative gene of each functionally homogeneous genomic cluster prior to enrichment analysis.
- For rare genetic variants, case-control pathway “burden” tests are the most appropriate pathway enrichment analysis method (**Box 3**).

### Future perspectives

Current pathway enrichment analysis methods provide a useful high-level overview of the pathways active in a genomics experiment. However, these methods consider a simplified pathway view (gene sets). Next generation pathway analysis methods will integrate more biological pathway details, build pathway models based on multiple types of genomics data measured across many samples, and consider positive and negative regulatory relationships in the data (**Box 5**). For instance, qualitative mathematical modeling parameterized with single cell RNA-seq data may enable accurate predictions of drug combinations capable of treating a given disease under study.

## PROTOCOL

### INTRODUCTION

This step-by-step protocol explains how to complete pathway enrichment analysis using g:Profiler (gene list) and GSEA (ranked gene list), followed by visualization and interpretation using Enrichment Map, as explained above in the text. The example data provided for the g:Profiler analysis is a list of genes with frequent somatic single nucleotide variants (SNVs) identified in The Cancer Genome Atlas (TCGA) exome sequencing data of 3,200 tumors of 12 types^25^. The example data provided for the GSEA analysis is a list of differentially expressed genes in two types of ovarian cancer defined by TCGA.

## MATERIALS

### Equipment

Hardware requirements:

- A recent personal computer with Internet access and at least 8GB of RAM. Note: 1GB of RAM is sufficient to run GSEA analysis but Cytoscape requires at least 8GB.

Software requirements:

- Assume that consumers (*C*(*t*), population of predators) are close to a chemical stimulus that is frequented by a population of bacteria that is predated *C*(*t*). In our example, the stimuli growths proportional to the time.
- A contemporary web browser (e.g. Chrome) for pathway enrichment analysis with g:Profiler (Protocol 1A).
- Java Standard Edition. Java is required to run GSEA and Cytoscape. It is available at http://java.oracle.com Version 8 or higher is required.
- GSEA desktop application for pathway enrichment analysis protocol 1B. Download the latest version of GSEA from http://www.broadinstitute.org/gsea/downloads.jsp. We recommend the javaGSEA desktop application. Free registration is required.
- Cytoscape desktop application is required for enrichment map visualization. The latest version of Cytoscape can be downloaded at http://www.cytoscape.org - Cytoscape version 3.5.1 or higher.
- The following Cytoscape apps must be installed within Cytoscape for enrichment map visualization. Go to **Apps App→ manager** (i.e., open the Apps menu and select the item “App manager”).

- Enrichment Map, version 3.0 or higher,
- Clustermaker2, version 0.9.5 or higher,
- WordCloud, version 3.1.0 or higher,
- AutoAnnotate, version 1.2.0 or higher.

Data requirements:

- We provide example files that are listed following the protocol. We recommend saving all files in a personal project folder before starting.
- A gene list or ranked gene list of interest.

- **Protocol 1A** with g:Profiler requires a list of genes, one per line in a text file or spreadsheet, ready to copy and paste into a web page.

- Example data for **Protocol 1A**: Genes with frequent somatic single nucleotide variants (SNVs) identified in The Cancer Genome Atlas (TCGA) exome sequencing data of 3,200 tumors of 12 types^25^. The authors used their MuSiC software to find 127 cancer driver genes that displayed higher than expected mutation frequencies (Ref^25^ Supplementary Table 4, column B) (**Supplementary_Table_1_Cancer drivers.txt**). Genes are ranked in decreasing order of significance (FDR *q*-value) and mutation frequency (not shown).

- - **Protocol 1B** with GSEA requires a RNK file with gene scores.
  - A RNK file is a two-column text file with gene IDs in the first column and gene scores in the second column. All (or most) genes in the genome need to have a score and the gene IDs need to match those used in the GMT file.
  - Example data for **Protocol 1B**: A list of differentially expressed genes in ovarian cancer from TCGA (**Supplementary_Table2_MesenvsImmuno_RNASeq_ranks.rn k**). This cohort was previously stratified into four molecular subtypes based on gene expression data, defined as differentiated, immunoreactive, mesenchymal and proliferative^44,45^. We compared the immunoreactive and mesenchymal subtypes to demonstrate the protocol. **Supplementary Protocol 1** shows how this file was created.

- Pathway gene set database
  - In **Protocol 1A**, g:Profiler maintains an up-to-date set of pathway gene sets from multiple sources and no further input from the user is required.
  - A database of pathway gene sets is required for **Protocol 1B** (GSEA). Human_GOBP_AllPathways_no_GO_iea_July_01_2017_symbol.gmt (**Supplementary_Table3_Human_GOBP_AllPathways_no_GO_iea_J uly_01_2017_symbol.gmt**) contains a database of pathway gene sets used for pathway enrichment analysis in the standard GMT format, downloaded from http://baderlab.org/GeneSets. This file contains pathways downloaded on July 1, 2017 from seven original pathway gene sets data sources: Gene Ontology^27^, Reactome^28^, Panther^21^, NetPath^30^, NCI^46^, MSigdb^47^, and HumanCyc^29^. The latest version of this file can be downloaded from http://baderlab.org/GeneSets.
  - A GMT file is a text file where every line represents a gene set of a single pathway. Each line includes a pathway ID, name and the list of associated genes in a tab-separated format.

### Equipment setup

- **Protocol 1A** uses web-based software and just requires a web browser.
- **Protocols 1B, 2 and 3** require installation of software on a local computer (see Equipment section, above).

## PROCEDURE

### Part 1A - Pathway enrichment analysis of a gene list using g:Profiler

1 Open the g:Profiler website at http://biit.cs.ut.ee/gprofiler/ (**Figure 2**).
2 Paste the gene list (copy list from **Supplementary_Table_1_Cancer drivers.txt**) into the “Query” field in top-left corner of the screen. The gene list can be space-separated or one per line. The organism for the analysis, *Homo sapiens,* is selected by default. The input list can contain a mix of gene and protein IDs, symbols and accession numbers. Duplicated and unrecognized IDs will be removed automatically, while ambiguous symbols can be refined in an interactive dialogue after submitting the query.
3 Check the box next to "Ordered query". This option treats the input as an ordered gene list and prioritizes genes with higher mutation enrichment scores at the beginning of the list.
4 Check the box next to "No electronic GO annotations". This option will discard less reliable Gene Ontology (GO) annotations (IEA - inferred from electronic annotation) that are not manually reviewed.
5 Set filters on gene annotation data using the legend on the right. We recommend that the first pathway enrichment analysis only includes biological processes (BP) of GO and molecular pathways of Reactome. Keep the two checkboxes checked and uncheck all other boxes in the legend.
6 Click on "Show Advanced Options" to set additional parameters.
7 Set the dropdown values of "Size of functional category" to 5 (‘min’) and 350 (‘max’). Large pathways are of limited interpretative value, while numerous small pathways decrease the statistical power because of excessive multiple testing.
8 Set the dropdown "Size of query/term intersection" to 3. The analysis will only consider more reliable pathways that have three or more genes in the input gene list.
9 Click "g:Profile!" to run the analysis. A graphical image will be shown with detected pathways from top to bottom and associated genes of the input list left to right. Resulting pathways are organized hierarchically into related groups. g:Profiler uses graphical output by default and switches to textual output when a large number of pathways is found. g:Profiler returns only statistically significant pathways with p-values adjusted for multiple testing correction using a custom pathway-focused procedure. By default, results with corrected q-value below 0.05 are reported.
10 Use the dropdown menu "Output type" and select the option "Generic Enrichment Map (TAB)". This file is required for visualizing pathway results with Cytoscape and Enrichment Map.
11 Click "g:Profile!" again to run the analysis with the updated parameters. The required link "Download data in Generic Enrichment Map (GEM) format" will appear under the g:Profiler interface. Download the file from the link and save it on your computer in your project folder. Example results are contained in **Supplementary_Table4_gprofiler_results.txt.**
12 Download the required GMT file by clicking on the link "name" at the bottom of the Advanced Options form. The GMT file is a compressed ZIP archive that contains all gene sets used by g:Profiler (e.g., gprofiler_hsapiens.NAME.gmt.zip). The gene set files are divided by data source. Download and uncompress the ZIP archive to your project folder. All required gene sets for this analysis will be in the file hsapiens.pathways.Name.gmt (**Supplementary_Table5_hsapiens.pathways.NAME.gmt**).
13 Proceed to **Protocol 2.**

**Figure 2.**
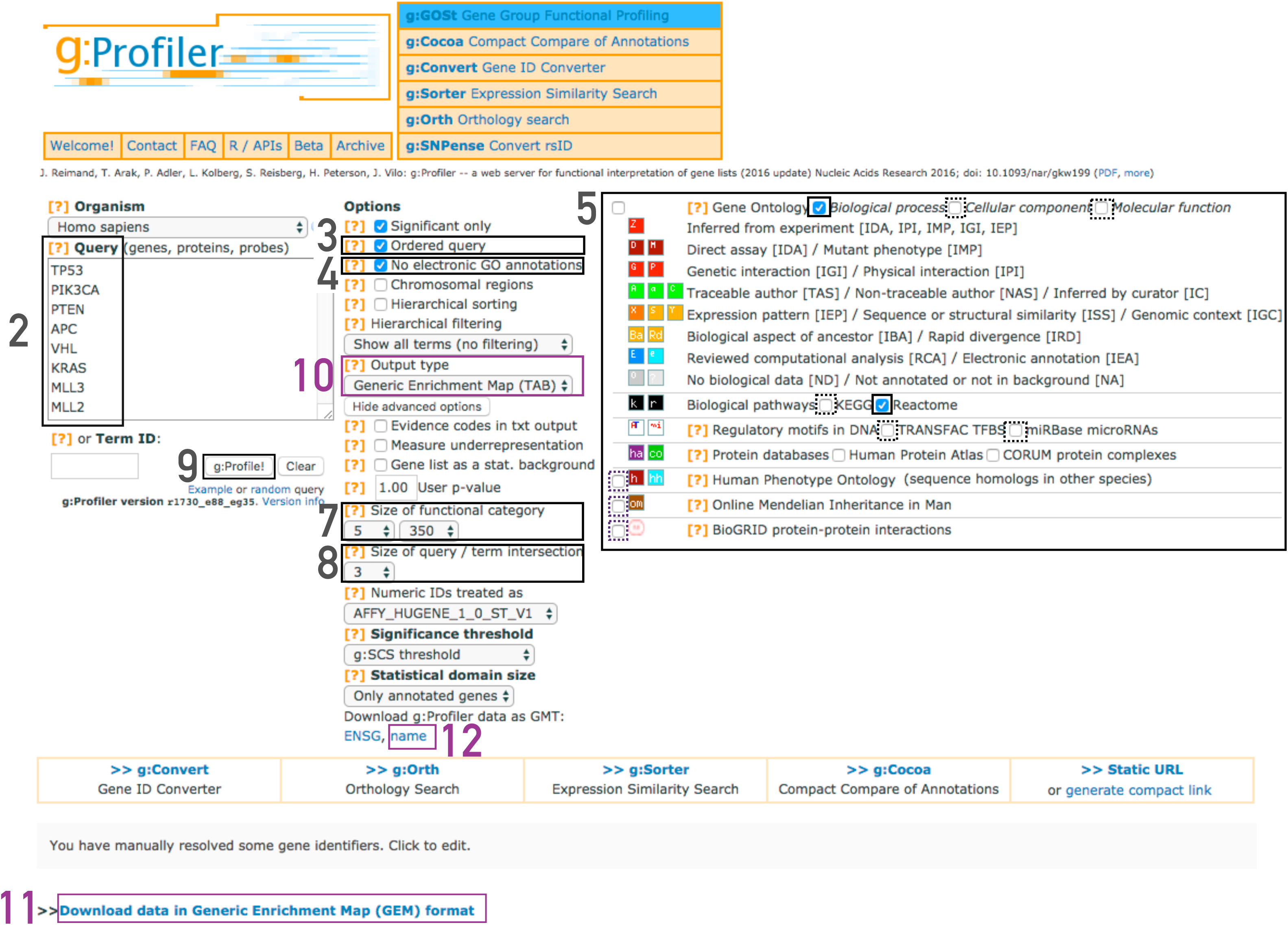
Screenshot of g:Profiler user interface. Steps in the protocol and associated g:Profiler interface components. Purple boxes indicate relate to files that need to be downloaded. The remaining boxes indicate parameters for the analysis.

#### TIMING

∼3 minutes to run g:Profiler using Chrome on Windows7.

### Part 1B - Pathway enrichment analysis of a ranked gene list using GSEA

14 Launch GSEA by double clicking on the downloaded GSEA file (gsea.jnlp) (**Tips and Troubleshooting – #1 (TT1)) (Figure 3**).
15 Load the required data files into GSEA:

i. Click on “Load Data” in the top left corner in the “Steps in GSEA Analysis” section.
ii. In the “Load Data” tab, click on “Browse for files …”
iii. Find your project folder and select the file Supplementary_Table2_MesenvsImmuno_RNASeq_ranks.rnk.
iv. Also select the pathway gene set definition (GMT) file using a multiple select method such as shift-click (**Supplementary_Table3_Human_GOBP_AllPathways_no_GO_iea_J uly_01_2017_symbol.gmt (TT2, TT3**)). Then click the ‘Choose’ button to continue.

**Figure 3.**
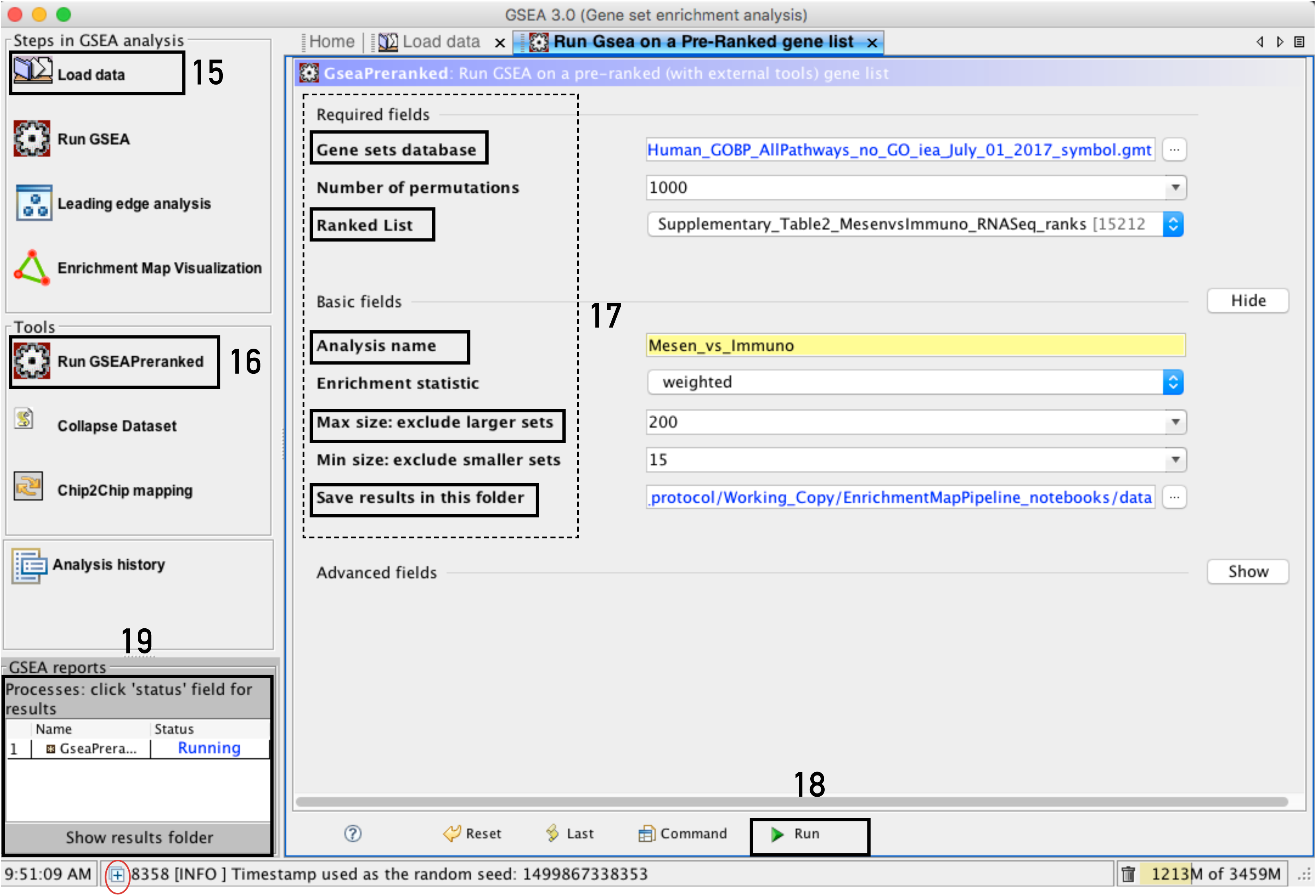
Screenshot of GSEA user interface. Steps in the protocol and associated GSEA interface components (version 3.0), highlighted with squares and with indication of the step number.

16 Click on “Run GSEAPreranked” in the side bar under “Tools”. The tab “Run GSEA on a Pre-Ranked gene list” will appear.
17 Specify the following parameters:

i. **Gene sets database** – click on the button (…) located to the right and wait for the gene set selection window to appear. Go to the “Gene matrix (local GMX/GMT)” tab using the top right arrow. Click on the downloaded local GMT file **Supplementary_Table3_Human_GOBP_AllPathways_no_GO_iea_Ju ly_01_2017_symbol.gmt** and click on OK at the bottom of the window.
ii. **Number of permutations** – number of times that the gene sets will be randomized to create the null distribution to calculate the p-value and FDR *q*-value (**TT4**). Use the default value of 1000 permutations.
iii. **Ranked List** – select the ranked gene list by clicking on the right-most arrow and highlighting the rank file (**Supplementary_Table2_MesenvsImmuno_RNASeq_ranks.rnk**).
iv. Click on “Show” button next to “Basic Fields” to display extra options.
v. **Analysis name** – change default “my_analysis” to a specific name, for example “Mesen_vs_Immuno”.
vi. **Save results in this folder** – navigate to the folder where GSEA should save the results. By default, GSEA will use gsea_home/output/[date] in your home directory.
vii. **Max size: exclude larger sets** – By default GSEA sets the upper limit to 500. Set this to 200 to remove the larger sets from the analysis.
18 Run GSEA - click on the “Run” button located at the bottom right corner of the window. Expand the window if the button is not visible. The “GSEA reports” panel at the bottom left of the window will show the status “Running”. It will be updated to “Success” upon completion (**TT5, TT6**).
19 Examine GSEA results - once the GSEA analysis is complete, a green notification “Success” will appear in the bottom left section of the screen. All output files are available in the folder specified in the GSEA interface. Click on “Success” to open the results in your web browser. Pathways enriched in top-ranking genes (i.e. up-regulated) are shown in the first set (na_pos; ‘mesenchymal’ in this protocol) and pathways enriched in bottom-ranked genes (i.e. down-regulated) in the second set (na_neg; ‘immunoreactive’) (**TT7, TT8**) (**Figure 4A**).

**Figure 4.**
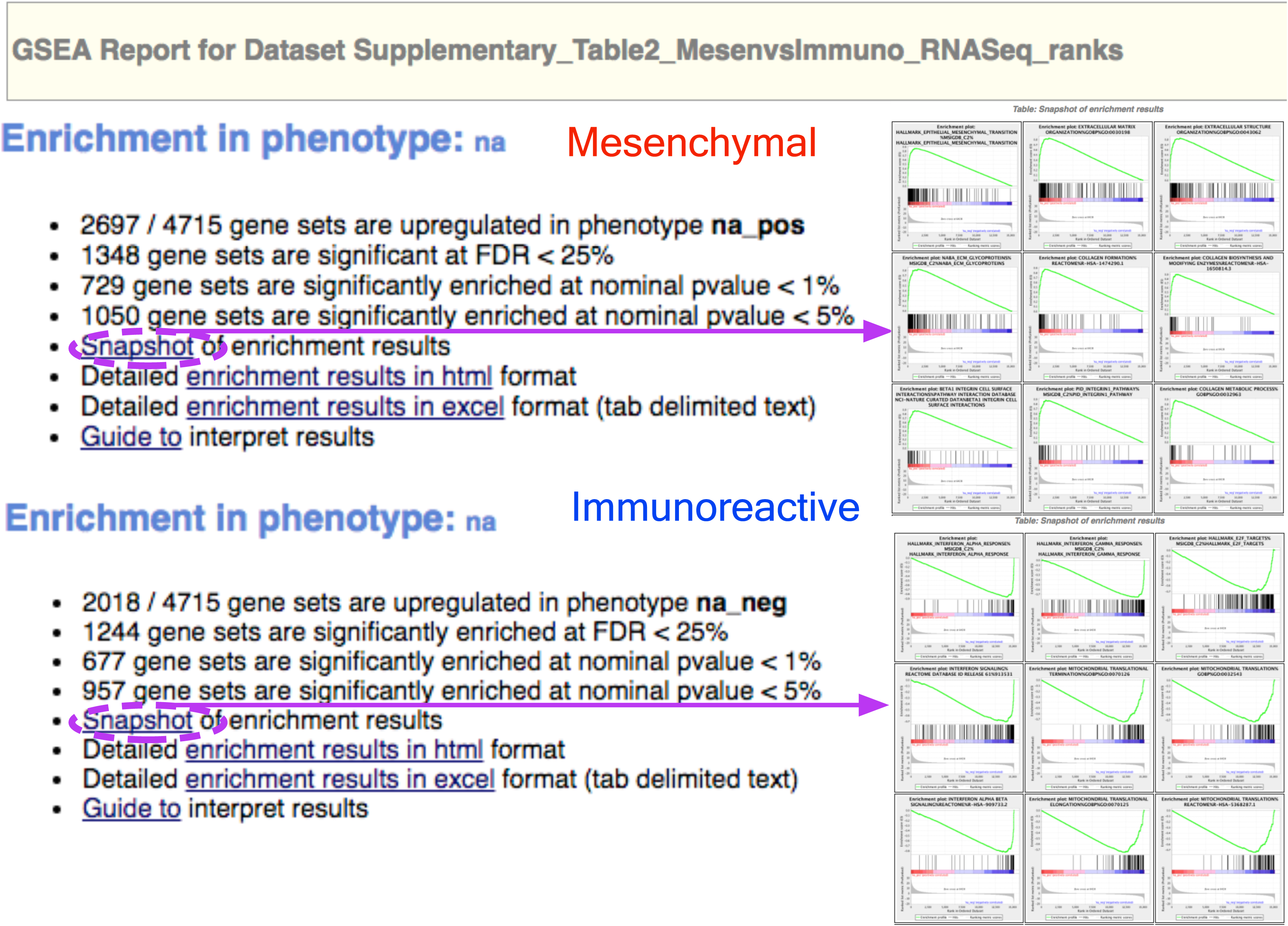
GSEA output. (A) GSEA output overview-Web page summary of GSEA results showing pathways enriched in the top or bottom of the ranked list, with “na_pos” and “na_neg” phenotype corresponding to enrichment in up-regulated and down-regulated genes. Clicking on snapshots under either of the phenotypes will show the top 20 enrichment plots for that phenotype. (B) Class/phenotype specific GSEA output in: the web page summary shows in purple how many gene sets were found enriched in up-regulated genes regardless of significance (purple), the total number of gene sets used after size filtering (cyan), the phenotype name (red), the number of gene sets that pass different thresholds (orange).

20 In the web browser results summary, click on the “Snapshot” link under the results to get an overview of the top 20 findings. The most significant pathways for the first phenotype (‘na_pos’) should clearly display enrichment in top-ranking (i.e. up-regulated) genes. Conversely, the most significant pathways for the second phenotype (‘na_neg’) should clearly display enrichment in bottom-ranked genes (i.e. down-regulated) (**TT9**) (**Figure 4A**).
21 Check the number of gene sets that have q-values below 0.05 to determine appropriate thresholds for the enrichment map in the next protocol. If no pathways are reported at *q*<0.05, more lenient thresholds such as *q*<0.1 or *q*<0.25 could be used (**Figure 4B**). The threshold *q*<0.25 provides very lenient filtering and it is not uncommon to find thousands of enriched pathways at this level. Robust analyses should use a cutoff of *q*<0.05 or lower. Filtering only by uncorrected p-values is not recommended.

#### TIMING

∼20 minutes to run GSEA using Windows7 with 1GB of RAM and Java 8 (TT4, TT5).

### Protocol 2 - Visualize enrichment results with Enrichment Map

22 Launch the Cytoscape software. Cytoscape introductory tutorials can be found at http://tutorials.cytoscape.org
23 In the menu, click Apps**→**Enrichment Map.
24 A "Create Enrichment Map Panel" will appear (**Figure 5**). Creating enrichment maps with g:Profiler and GSEA requires slightly different input files.

**Figure 5.**
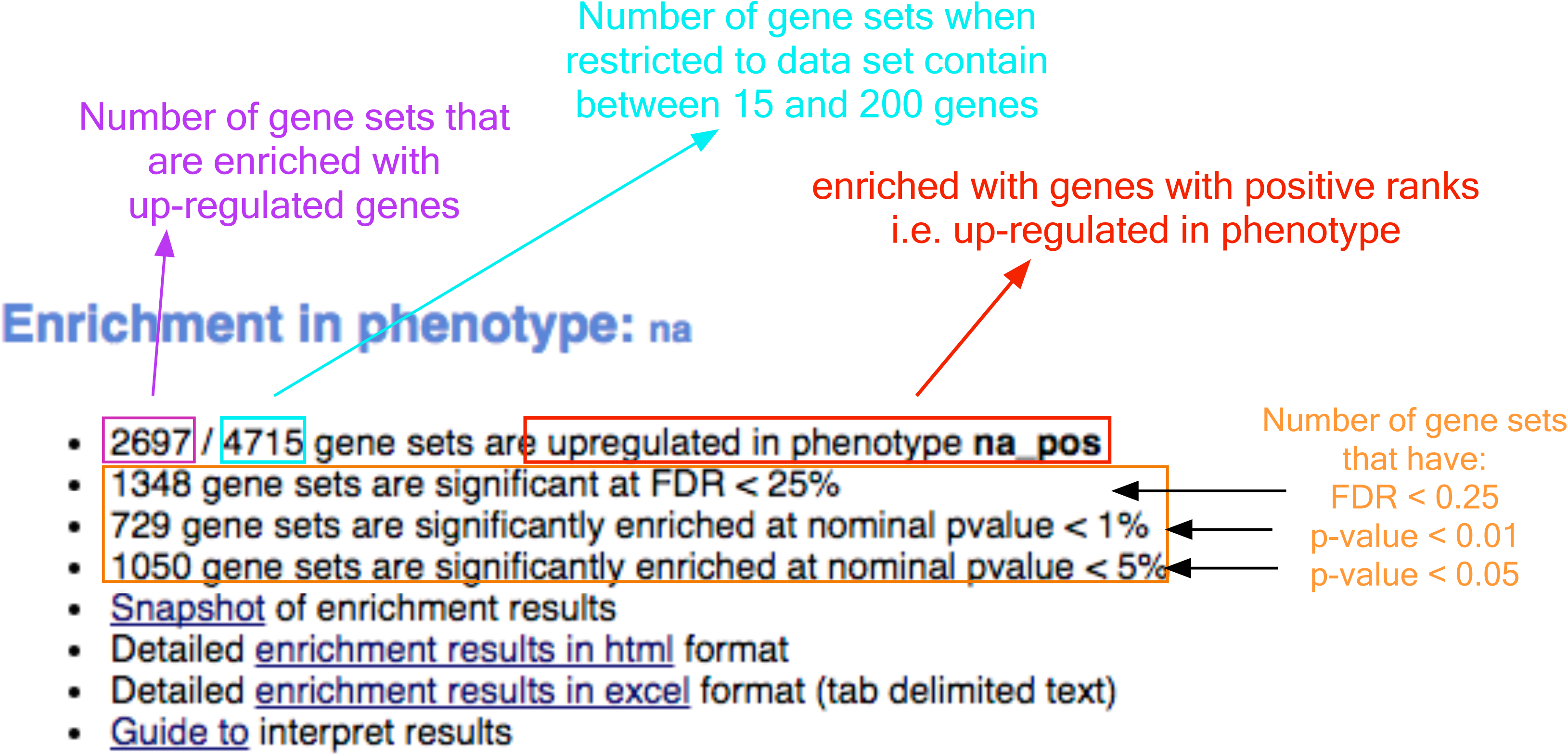
Screenshot of Enrichment Map user interface. Input fields in the Enrichment Map interface for (A) g:Profiler and (B) GSEA results. Other than the specific input files, the parameters are the same for both analysis types. Attributes surrounded by a dashed box should be filled out automatically if the user selects a folder for enrichment map to find all analyses in. If the file names are missing, then EM was unable to find the specified file, possibly because the original file was moved or deleted. Orange boxes indicate optional files. For the examples presented in the protocol, optional files are used for the GSEA analysis but not for the g:Profiler analysis to demonstrate the two distinct use cases.

25 For g:Profiler results, files generated in **Protocol 1A** (**Figure 5A**):
  i. Click on folder icon to navigate to the g:Profiler results
  ii. Click on g:Profiler folder created in Protocol 1A. Click on Open. (**TT10**)
  iii. In the right hand panel g:Profiler output files will be auto populated into their specified fields. (Alternately, users can click on the “+” to specify each of the required files manually).

- If desired, modify the ‘dataset’ name. By default, EM will use the name of the g:Profiler enrichment results file name (e.g. gprofiler_cancer_drivers).
- Verify the Analysis type is set to “Generic/gProfiler”.
- Verify the Enrichments results file is the results file downloaded in Protocol 1A step 11 (or alternately manually specify **Supplementary_Table4_gprofiler_results.txt**)
- Verify the GMT specified is the file retrieved from the g:Profiler website in Protocol 1A – step 12. Use the file hsapiens.pathways.NAME.gmt (or alternately manually specify **Supplementary_Table5_hsapiens.pathways.NAME.gmt**) that contains gene sets corresponding to GO biological processes and Reactome pathways.
  iv. Specify additional files:

- **Expression** - (Optional) Upload an expression matrix for the genes analyzed in g:Profiler or alternatively an expression data set of all genes. If the expression data set contains additional genes not used for the g:Profiler search, their expression values will still appear in the heat map of the enrichment map (for example file see **Supplementary_Table6_TCGA_OV_RNAseq_expression.txt**).
- **Ranks** – (Optional) Ranks for gene list or for the expression data can also be specified (for example file see **Supplementary_Table2_MesenvsImmuno_RNASeq_ranks.rnk**
- **Classes** – (Optional) GSEA cls file defining the phenotype (i.e. biological conditions) of each sample in the expression file, for example file see **Supplementary_Table9_TCGA_OV_RNAseq_classes.cls.** Generally, this file is only required when performing phenotype randomization in GSEA, but if it is supplied to enrichment map it is used to identify and label the columns of the expression file in the Enrichment Map heat map by phenotype.
- **Phenotypes** – (Optional) If there are two different phenotypes in the expression data, update the phenotype labels so that ‘positive’ represents the phenotype associated with positive values (Mesenchymal in this example) and ‘negative’ with negative values (Immunoreactive in this example) (**TT11**). Tune parameters in the “Parameters” box:

- g:Profiler automatically retrieves only statistically significant results (*q*<0.05), so the q-value and P-value cutoff parameters can be set in the Enrichment Map Input panel to 1, unless more stringent filtering is desired. For this analysis set FDR *q*-value to 0.01.
- Keep the connectivity slider in the center. If the network is too cluttered because it has too many connections (edges), move the slider to the left to make the network sparser. Alternatively, if the network is too sparse (i.e. there are too many disconnected pathways), move the slider to the right to obtain a more densely connected network (**TT12**).
- Click the “Build” button at the bottom of the Enrichment Map Input panel.
26 For GSEA results generated in **Protocol 1B (Figure 5)**:

i. Click on folder icon to navigate to the GSEA results
ii. Click on GSEA folder created in Protocol 1B. Click on Open. (**TT13**)
iii. In the right-hand panel, GSEA output files will be auto populated into their specified fields. Alternately the “+” can be clicked to specify each of the required files manually. Equivalent supplementary files that users can specify manually are indicated in brackets.

- If desired, modify the ‘dataset’ name. By default, EM will use the name of the GSEA results folder prior to the first ‘.’ as the ‘dataset’ name.
- Verify the Analysis type is set to “GSEA”.
- **GMT** – Verify that the file is set to [data-directory]/**Supplementary_Table3_Human_GOBP_AllPathway s_no_GO_iea_July_01_2017_symbol.gmt** (or alternately navigate to the following file: **Supplementary_Table3_Human_GOBP_AllPathways_no_GO _iea_July_01_2017_symbol.gmt)(TT14, TT15**)
- **Enrichments 1** – Verify that the file is set to [path_to_gsea_dir]/Mesen_vs_Immuno.GseaPreranked. [unique number]/gsea_report_for_na_pos_[unique number].xls where [unique number] is a number generated by GSEA (see TT14) and path_to_gsea_dir is the full path to the directory selected in step 26ii, above, or alternately navigate to the following file: **Supplementary_Table7_gsea_report_for_na_pos.xls**
- **Enrichments 2** – Verify that the file is set to [path_to_gsea_dir]/Mesen_vs_Immuno.GseaPreranked. [unique number]/gsea_report_for_na_neg_[unique number].xls where path_to_gsea_dir is the full path to the directory selected in step 26ii, above, or alternately navigate to the file: **Supplementary_Table8_gsea_report_for_na_neg.xls (TT14)**
- **Ranks** – Verify that the file is set to [path_to_gsea_dir]/Mesen_vs_Immuno.GseaPreranked.[unique number]/ranked_gene_list_na_pos_versus_na_neg_[unique number].xls where path_to_gsea_dir is the full path to the directory selected in step 26ii, above, or alternately navigate to the file: **Supplementary_Table2_MesenvsImmuno_RNASeq_ranks.rnk**
iv. Specify additional files:

- **Expression** – (Optional) Supplementary_Table6_TCGA_OV_RNAseq_expression.txt
- **Classes** – (Optional) Supplementary_Table9_TCGA_OV_RNAseq_classes.cls
- **Phenotypes** – (Optional) In the text boxes replace ‘na_pos’ with "Mesenchymal" and ‘na_neg’ with Immunoreactive. Mesenchymal will be associated with red nodes as it corresponds to the positive phenotype while Immunoreactive will be labeled blue (**TT16, TT17**).
v. Tune parameters in the “Parameters” box:

- Set FDR q-value cutoff to 0.01 (**TT18**).
- Keep the connectivity slider in the center. For networks with fewer edges, a sparser network, move the slider to the left. Alternatively, for networks with more edges, a denser network, move the slider to the right (**TT12**).
vi. Click the “Build” button at the bottom of the Enrichment Map Input panel (**TT6**).
27 **Figure 6** shows the resulting enrichment maps from the above g:Profiler and GSEA protocols.

**Figure 6.**
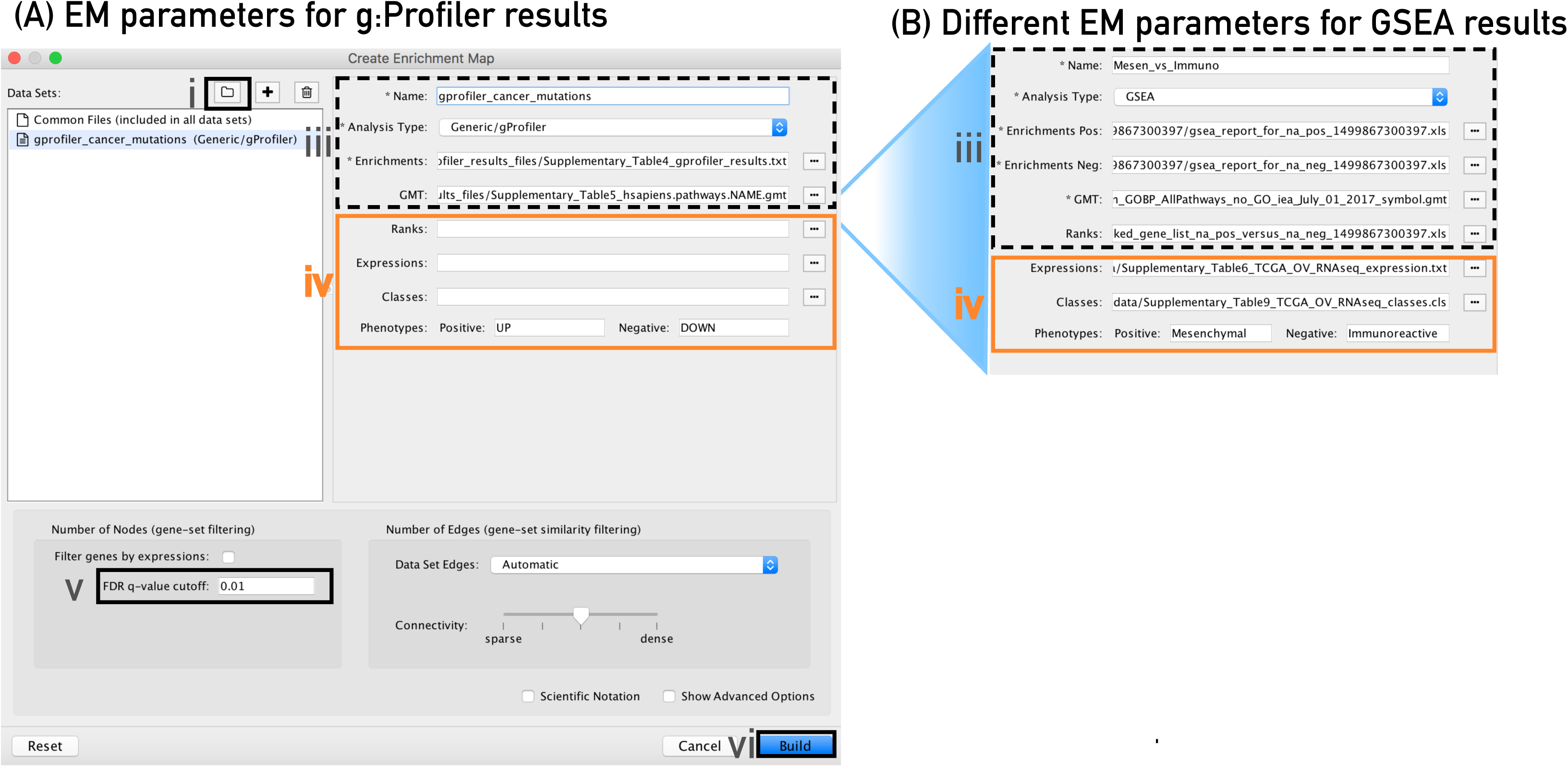
Resulting Enrichment Maps (no manual formatting) Unformatted enrichment maps generated from Protocols 1A and 1B. Each node (circle) represents a distinct pathway and edges (blue lines) represent the number of genes overlapping between two pathways determined using the similarity coefficient. (A) Enrichment map of significantly mutated cancer driver genes generated using the g:Profiler analysis in Protocol 1A. (B) Enrichment map of pathways enriched in up regulated genes in Mesenchymal (red) and Immunoreactive (blue) ovarian cancer samples using the GSEA analysis in Protocol 1B.

### TIMING

∼5 minutes to create Enrichment Map in Cytoscape using Windows7 with 8GB of RAM and Java 8.

### Protocol 3 - Navigating and interpreting the Enrichment Map

An enrichment map must be interpreted to discover novel information about a set of data and must be refined to create a publication quality figure.

28 To explore the enrichment map, select the network of interest in the control panel located at the left side of the Cytoscape window and navigate it (zoom and pan) using Cytoscape controls (**Figure7A**). Pathways with many common genes often represent similar biological processes and group together as ‘themes’ in the network. Click on a node to display the corresponding genes in the table below the network view (**Figure 7B**).

**Figure 7:**
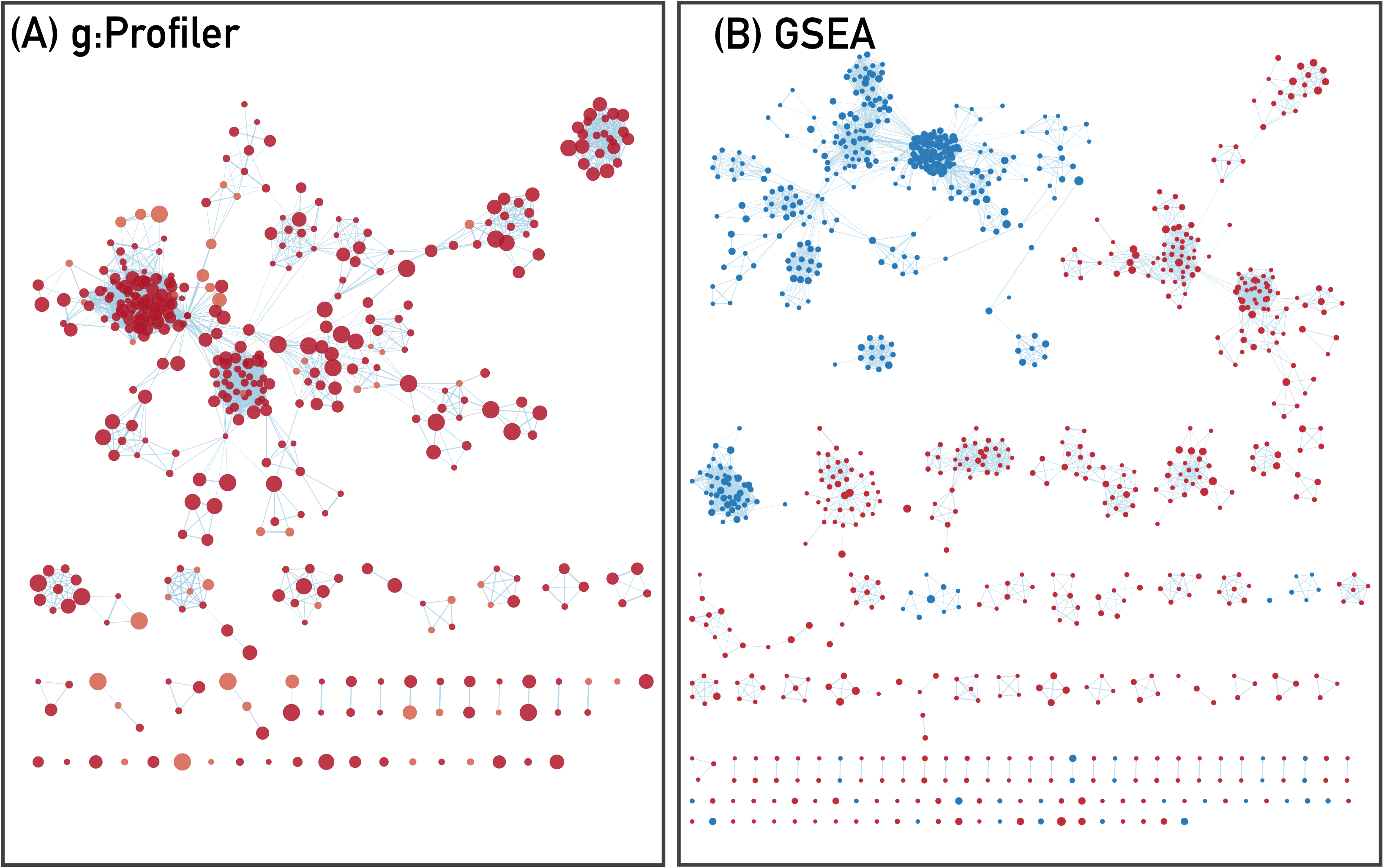
Overview of Cytoscape panels. (A) Cytoscape control panel that contains networks, styles and select panels as well as the Enrichment Map main panel. (B) The table panel contains tables with node, edge and network attributes. An Enrichment map heat map panel displaying expression for genes associated with selected nodes and edges is available in this panel. (C) Cytoscape search bar. (D) Node table containing values for all variables associated with each node in the network.

29 To find a gene or pathway of interest, type its name in the search bar located in the top right corner (**Figure 7C**). All pathways containing that gene will be highlighted. For example, *TP53* and *BGN* are the top genes in g:Profiler and GSEA analyses, respectively (**TT19**).
30 To find the most enriched pathways, find the column named “EM1_fdr_qvalue” (for g:Profiler) or EM1_NES” (for GSEA) in the ‘Node’ tab in the table panel (**Figure 7C and 7D**). For GSEA, we specifically recommend using the NES (normalized enrichment score) to sort pathways by enrichment strength, whereas we recommend using the enrichment *p*-value for other enrichment methods (**TT21**). Click on the column name to sort the table according to that attribute. Click the greatest value to show the pathway most enriched in the data. To highlight a subset of the pathways in the network, select pathways of interest, right-click on any selected row in the table and select “Select nodes from selected rows”.
31 When a gene expression matrix is provided as input to enrichment map (**TT21**), we can study the enriched pathway gene expression patterns. Click on an individual node to generate a gene expression heat map in the table panel (**Figure 8**). If the analysis is based on GSEA results and a rank file is supplied, the leading-edge genes are highlighted in yellow for individual node selections (**TT22**). To improve the heat map visualization, in the table panel, “Heat Map” tab, change:

i. Adjust the Sorting options (**Figure 8A**) - by default the heat map is sorted by ranks if a rank file is supplied. In the absence of a rank file no sort is applied. Sorting options include hierarchical clustering, ranks, or no sorting (**Figure 8F**). Additional rank files can be uploaded through the settings menu in the heat map panel for comparison. Clicking on any of the column names will change the sorting to the selected column. Clicking on the arrow next to the currently sorted column will invert the order of sorting (**TT23**).
ii. Define genes you wish to include in the heat map (**Figure 8B**) – data can be viewed for all genes contained in the selected nodes or just for the genes common to all selected nodes. By default all genes are shown.
iii. Change expression value visualization depending on your data type (**Figure 8C**) – data can be viewed as it was loaded (“Values”), or row normalized where the row mean is subtracted from every value and then divided by the row’s standard deviation (“Row Norm”), or log transformed (“log”).
iv. Change Expression viewing options (**Figure 8D**) – By default, all expression values are visible in the heat map for expression sets with 49 or fewer samples. Above that, EM will automatically compress the values to their median value. Other options include no compression (“-None-”), minimum values (“Min”), or maximum value (“Max”).
v. To see the individual expression values, select “show values” (**Figure 8E**).
vi. Additional fine tuning of the heat map can be done through the settings panel that includes functionality to add new rank files, export the heat map data as a tab delimited text file or PDF image, change the distance metric for hierarchical clustering, or turn on heat map autofocus (**Figure 8F**). The resulting heat map can be seen in **Figure 8**. Columns headings are colored according to sample phenotype. Red color refers to the first phenotype (Mesenchymal), and blue to the second phenotype (Immunoreactive) (TT24). The heat map can be exported to a text file for further analysis.

- Click on “Export to txt” in heat map settings (**Figure 8F**)
- Specify the name and location of the saved file
- If only an individual node is selected, a dialog will offer to save the leading edge only. If “Yes”, only the highlighted genes will be exported, and the entire set is exported otherwise (**TT25**)

**Figure 8:**
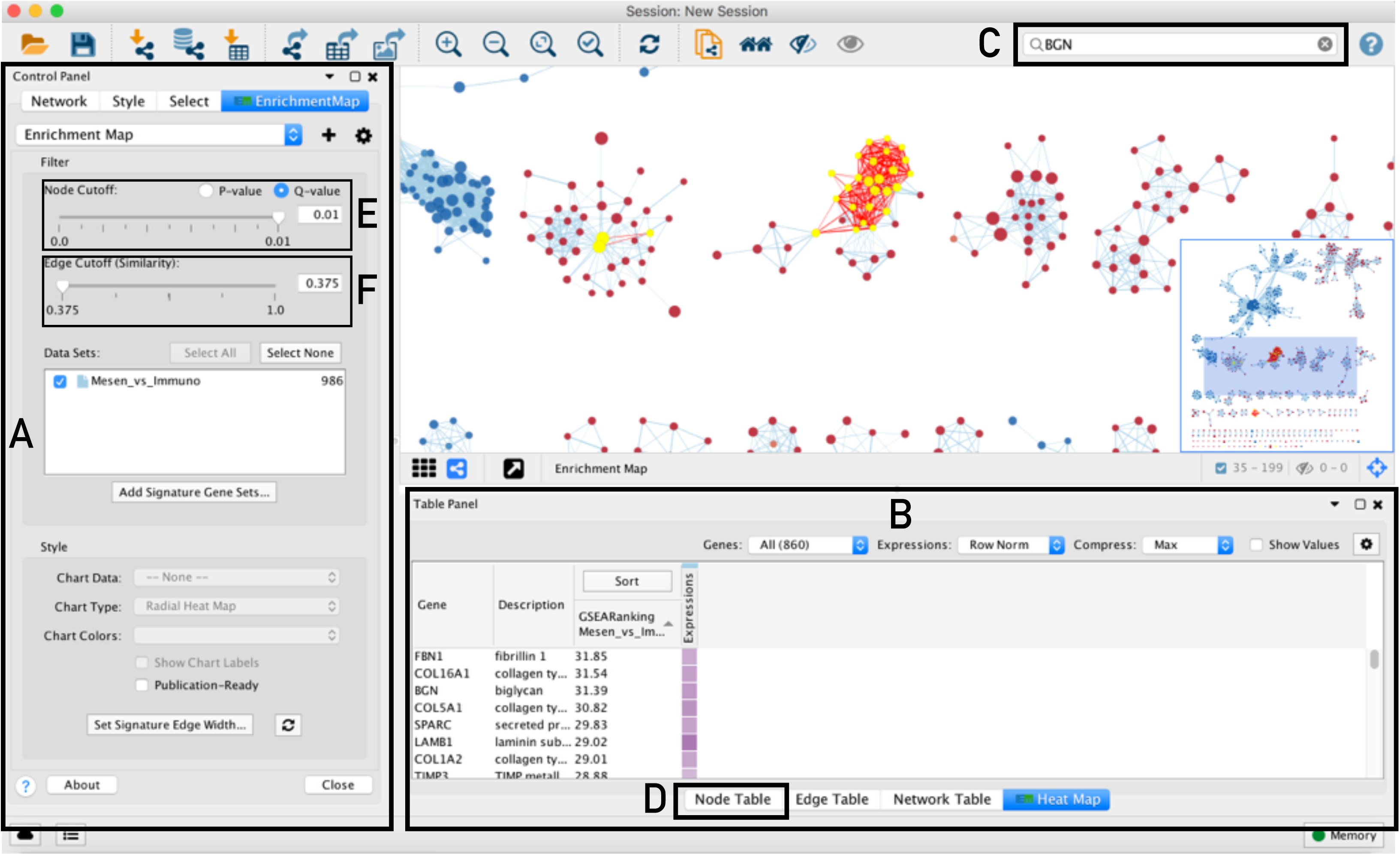
Example Enrichment Map heat map. Heat map created by selecting the Immunoreactive pathway interferon alpha beta signaling pathway from Reactome. This heat map is for GSEA results, thus the leading edge genes are highlighted in yellow. Additional controls in the heat map panel include: (A) sorting options, (B) genes to include, (C) expression data visualization options, (D) data compression options, (E) to show values, and (F) adjust settings.

32 Organize and de-clutter the network

i. If the network has too many nodes, increasing the Node cutoff *q*-value will remove less significant nodes (**Figure 7E**).
ii. If the network is too interconnected, increasing the edge cutoff (similarity) threshold will remove less pronounced edges between nodes (**Figure 7F**).
iii. The network layout may be applied again after adjusting the cutoffs (see the Layout menu in Cytoscape). The default layout algorithm is the unweighted prefuse force-directed layout. We also recommend the yFiles organic layout or weighted prefuse force-directed layouts. (**TT26**)
iv. To restore nodes or edges, adjust threshold sliders to their original positions.
v. It can be helpful to separate the two different phenotypes (i.e. place all the red nodes to one side and all blue nodes to the other). To do this:

- Click on the select tab in the control panel (**Figure 7A**)
- Click on the “+” and select “Column filter”
- Click on “Choose column.” and select “EM1_NES (Mesem_vs_Immuno)”
- Click on the box next to “between” and change the value to zero. Click “Enter”
- All red nodes should now be selected. Click and hold on any selected node and drag selection to the left until it does not overlap any blue nodes
- Click on Layouts menu. Select “Prefuse Force Directed Layout” --> “Selected Node Only”**→**“(none)” (**TT26**)
- In the Select tab, adjust slider to select all negative values. Click on “Apply” at the bottom of the Select tab
- All blue nodes should now be selected. Click and hold on any selected node and drag selection to the right until it does not overlap any red nodes
- Click on Layouts menu. Select “Prefuse Force Directed Layout” --> “Selected Node Only”**→**“(none)” (**TT26**)
33 Define major themes. Enrichment maps typically include clusters of similar pathways representing major biological themes. Clusters can be automatically defined and summarized using the AutoAnnotate Cytoscape app. AutoAnnotate first clusters the network using the clusterMaker app and then summarizes each cluster based on word frequency within the pathway names via the WordCloud app (**TT27, TT28**).

i. Launch AutoAnnotate by selecting Apps**→**AutoAnnotate**→**New Annotation Set. in the Cytoscape menu bar. The “AutoAnnotate” tab will appear in the Cytoscape control panel.
ii. Click on “+” in the AutoAnnotate panel.
iii. The “AutoAnnotate: create Annotation Set” panel will appear.
iv. In the “Quick Start” tab click on “Create Annotations” (**TT29**).
v. Each cluster in the network will have a circle annotation drawn around it and will be associated with a set of words (by default three) that appear most in the node description fields. Moving individual nodes within a cluster will automatically resize the surrounding circle annotation and moving an entire cluster will redraw the annotations in the new cluster location (**TT30**).
vi. Manually arrange clusters to clean up the figure. Move nodes to reduce node and label overlap. **Figure 9** shows the results of this process.

**Figure 9.**
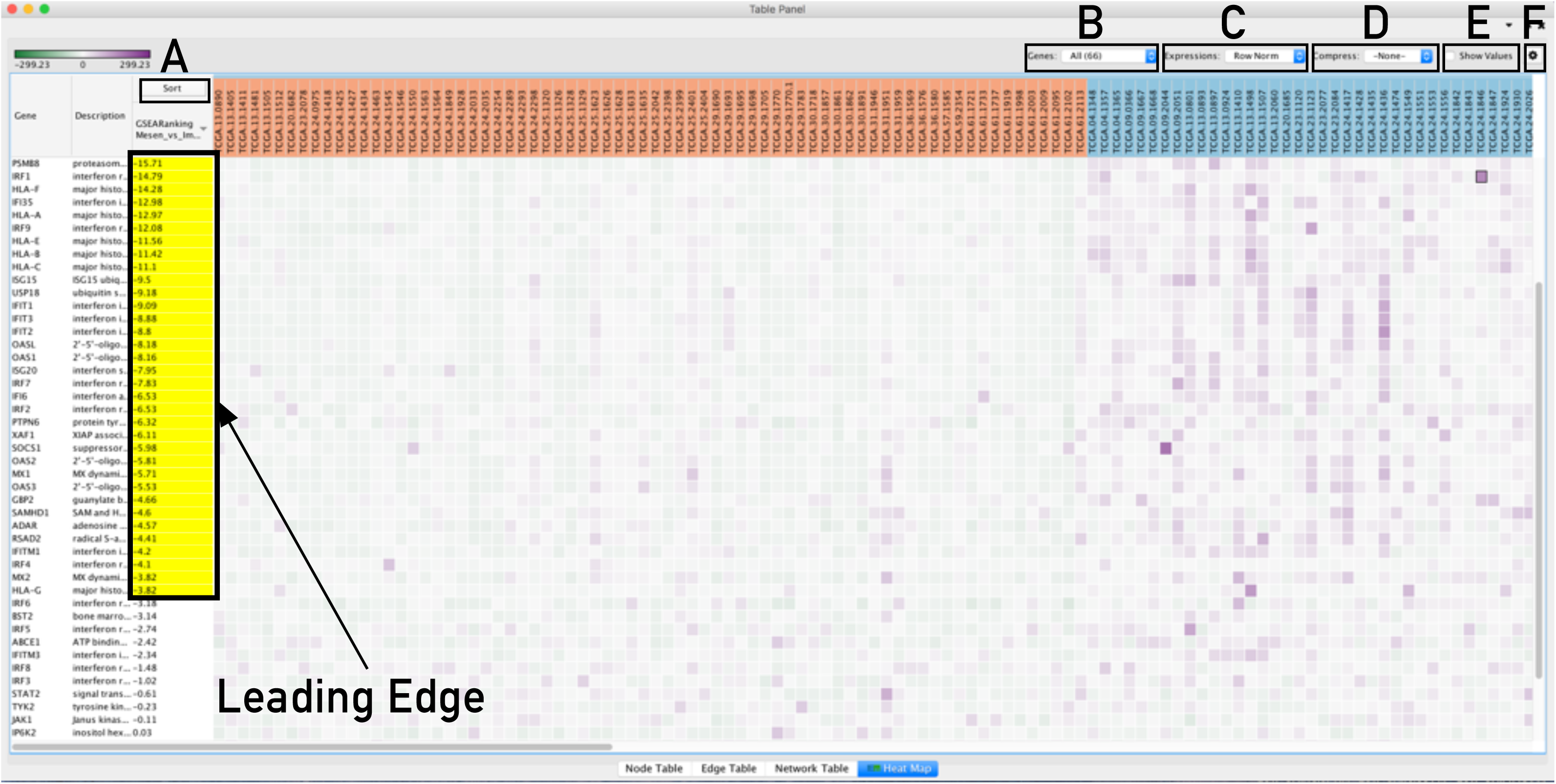
Resulting Publication Ready Enrichment Map. Publication-ready annotated enrichment map (created with parameters FDR *q*-value < 0. 01, and combined coefficient >0.375 with combined constant = 0.5). Red and blue nodes represent mesenchymal and immunoreactive phenotype pathways, respectively, and were manually separated to form a clearer picture. Clusters of nodes were labelled using the AutoAnnotate Cytoscape app. Individual node labels were removed for clarity using the publication ready button in Enrichment Map and exported to PNG and PDF files. The figure was resized using illustration software but no additional modifications were made to the network.

34 Create a simplified network view (**Figure 10**). This creates a single group node for every cluster with a summarized name and provides an overview of the enrichment result themes that is useful for enrichment maps containing many nodes.
  1. In the Cytoscape Control Panel select the “AutoAnnotate” Tab.
  2. Click on the Menu icon in the upper right hand corner.
  3. Select “Collapse All” (**TT31, TT32, TT33**).
  4. Scale collapsed network for better viewing:

i. In the Cytoscape menu bar, select: View → Show Tool Panel.
ii. Go to Tool Panel located at the bottom of the Control Panel.
iii. Click on the “Scale” Tab.
iv. Move slider left to tighten the node spacing.

**Figure 10.**
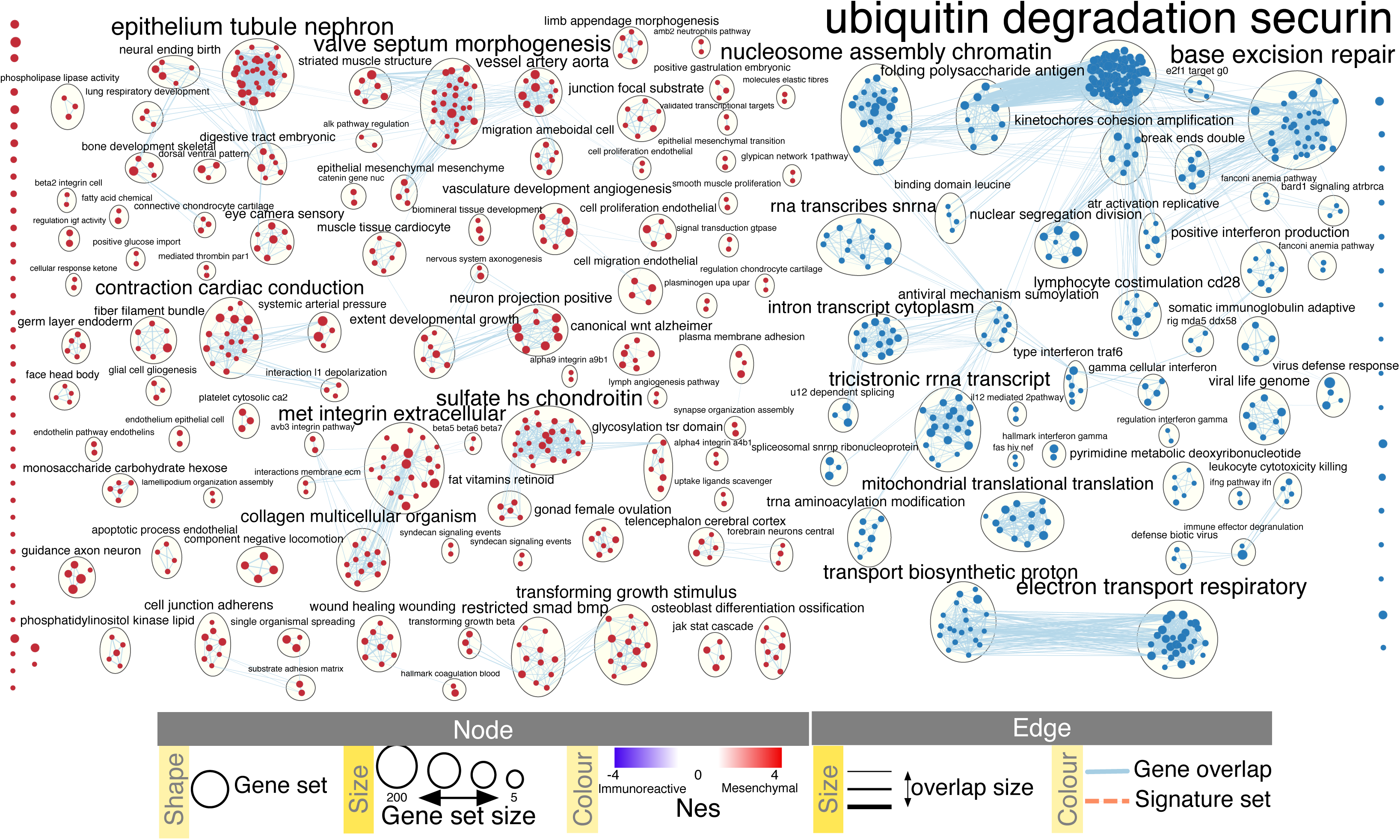
Collapsed Enrichment Map. The network was further summarized by collapsing node clusters using the AutoAnnotate app. The network was scaled for better node distribution and manually adjusted to reduce node and label overlap. The enrichment map was exported to PNG and PDF. No additional modifications were made to the network in any illustration software tools.

35 Manually arranging the network nodes and custom labeling the major themes is required for the clearest network view and for a publication quality figure.

i. For instance, it is useful to bring together similar themes, such as signaling or metabolic pathways, even if they are not connected in the map.
ii. If the focus of the figure is only on a subset of the network, it can be easier to work with just the subset. To create this, select the nodes of interest, then in the Cytoscape menu bar Select File**→**New**→**Network**→**From selected nodes, all edges.
iii. When the purpose of the figure is to show a large network highlight only the main themes, clicking on “Publication ready” in the enrichment map panel will remove node labels. To revert to the original network, click on the “Publication ready” button again.
36 Create a sub network that highlights a specific theme or data – often enrichment maps generated from platforms that measure signals from a large percentage of the genome are large and complicated. When generating a figure, it is important to highlight specific themes or pathways relevant to the analysis in question. For example, we will select the top mesenchymal and immunoreactive pathways and create a sub network containing them.

i. Click on the select tab in the control panel (**Figure 7A**).
ii. Click on the “+” and select “Column filter”.
iii. Click on “Choose column.” and select “EM1_NES (Mesem_vs_Immuno)”.
iv. Click on the box next to “between” and change the value to 2.5. Click “Enter”.
v. Click on the “+” and select “Column filter”.
vi. Click on “Choose column.” and select “EM1_NES (Mesem_vs_Immuno)”.
vii. Click on the box next to “inclusive” and change the value to -2.5. Click “Enter”.
viii. Above the two column filters you just added, change the drop down from “Match all (AND)” to “Match any (OR)”.
ix. Click on Apply. Under the apply button, it should say “Selected 32 nodes and 0 edges in Xms”. The exact number of seconds specified will depend on your computer speed.
x. Select File**→**New**→**Network**→**From selected nodes, all edges.
xi. A new smaller network should appear. Manually move nodes around for optimal layout.
xii. Annotate network as described in step 6 (Figure 11).

**Figure 11.**
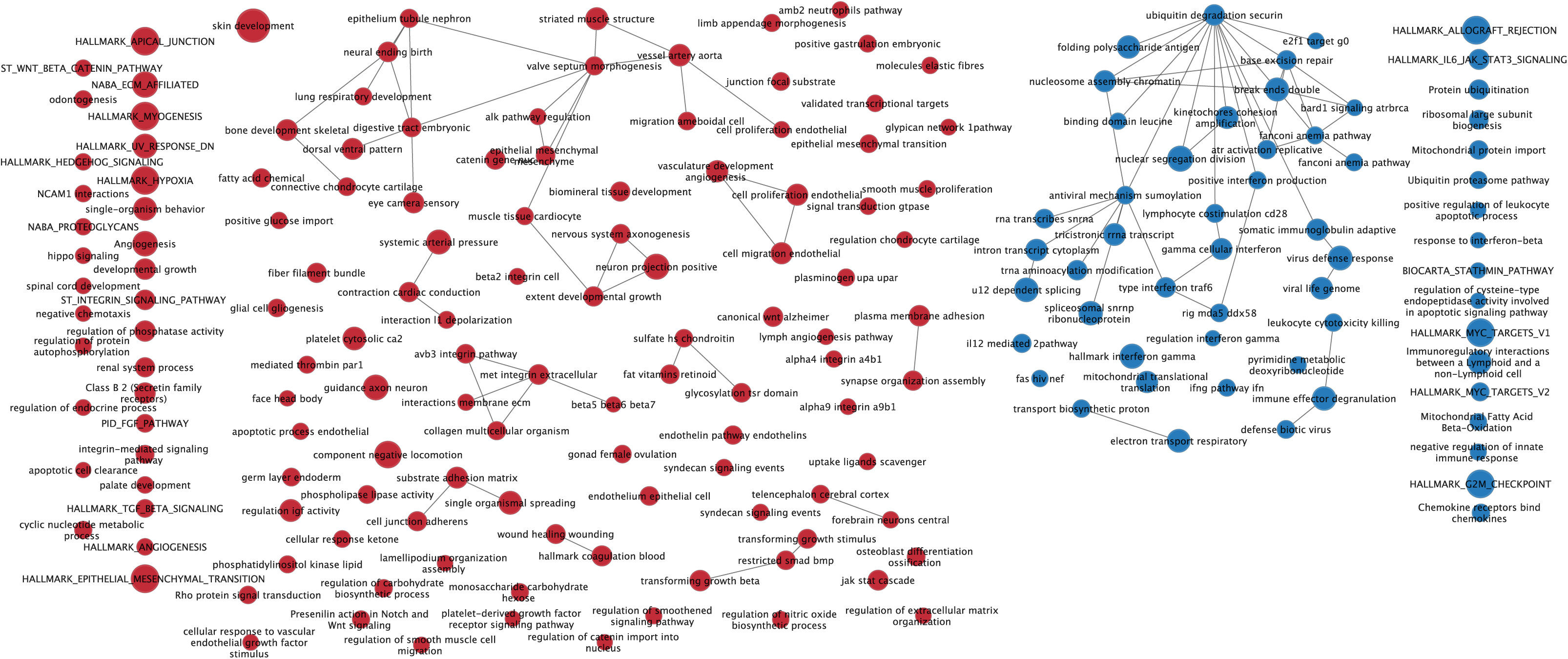
Subnetwork Example. Sub network of the main Enrichment Map figure (Figure 9) manually created by selecting pathways with the top NES values and creating a new network from the selection. Red and blue nodes are mesenchymal and immunoreactive phenotypes, respectively. Clusters of nodes were automatically labelled using the AutoAnnotate Cytoscape app. Annotations in the subnetwork may differ slightly from the main network as word counts were normalized on a network basis.

37 Export the image (**TT34**)

i. In the Cytoscape menu bar, select File → Export as Image…
ii. Set “Select the export file format” to PDF (**TT35**).
iii. Click on “Browse.” to specify file name and location.
iv. Click on “Save” to close the browser window and then on “OK”.
v. A window “Export Network” will appear, click on the “OK” button.
38 Get network creation parameters. In the previous step we exported the network as an image but there is information that either needs to be included in the text legend or as a pictograph within the image itself so the network can be easily interpreted. It is important to include the thresholds used when creating the map.

i. In the Enrichment Map Input panel click on the cog (settings) icon in the top right hand corner.
ii. Click on show legend.
iii. In the EnrichmentMap Legend panel click on the “Creation Parameters…”
iv. In the displayed panel you will find the thresholds to be added to the figure legend. Add FDR *q*-value, similarity metric and threshold to text legend of figure. For example: “Enrichment map was created with parameters *q* < 0.01, and combined coefficient >0.375 with combined constant = 0.5 (**TT36**)”.
39 Create a legend - there are many different node and edge attributes used in the enrichment map to represent different aspects of the data. It is important to add their meaning in the text legend or as a pictograph in the figure. Although Cytoscape has the ability to export a legend of the current style, it is not easily transferrable as a legend for the resulting figure. **Figure 12** shows the basic legend components (available as SVG and PDF images at http://baderlab.org/Software/EnrichmentMap#Legends)) that can be used for an enrichment map figure. Only include components relevant to the given analysis. See bottom of **Figure 9** for components used for current analysis.

**Figure 12.**
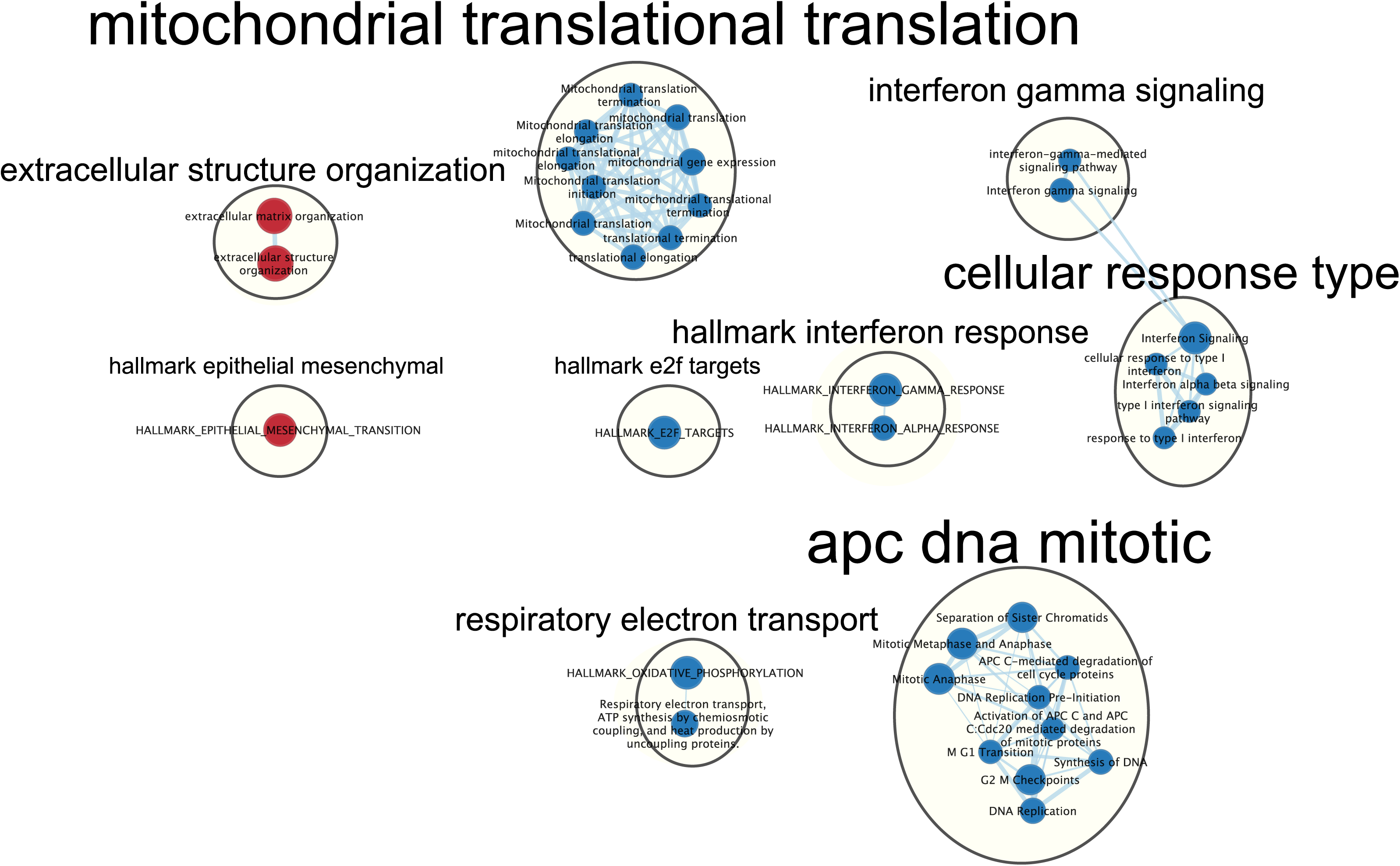
Generic enrichment map legend. Enrichment map attributes that can be copied for use in a figure legend. Only include components relevant to the analysis depicted in the figure. Post analysis ‘Signature set’ nodes are included in the generic legend even though they are not covered in this protocol. Post analysis nodes are used to highlight pathways in the enrichment map that contain genes of interest such as drug or microRNA targets.

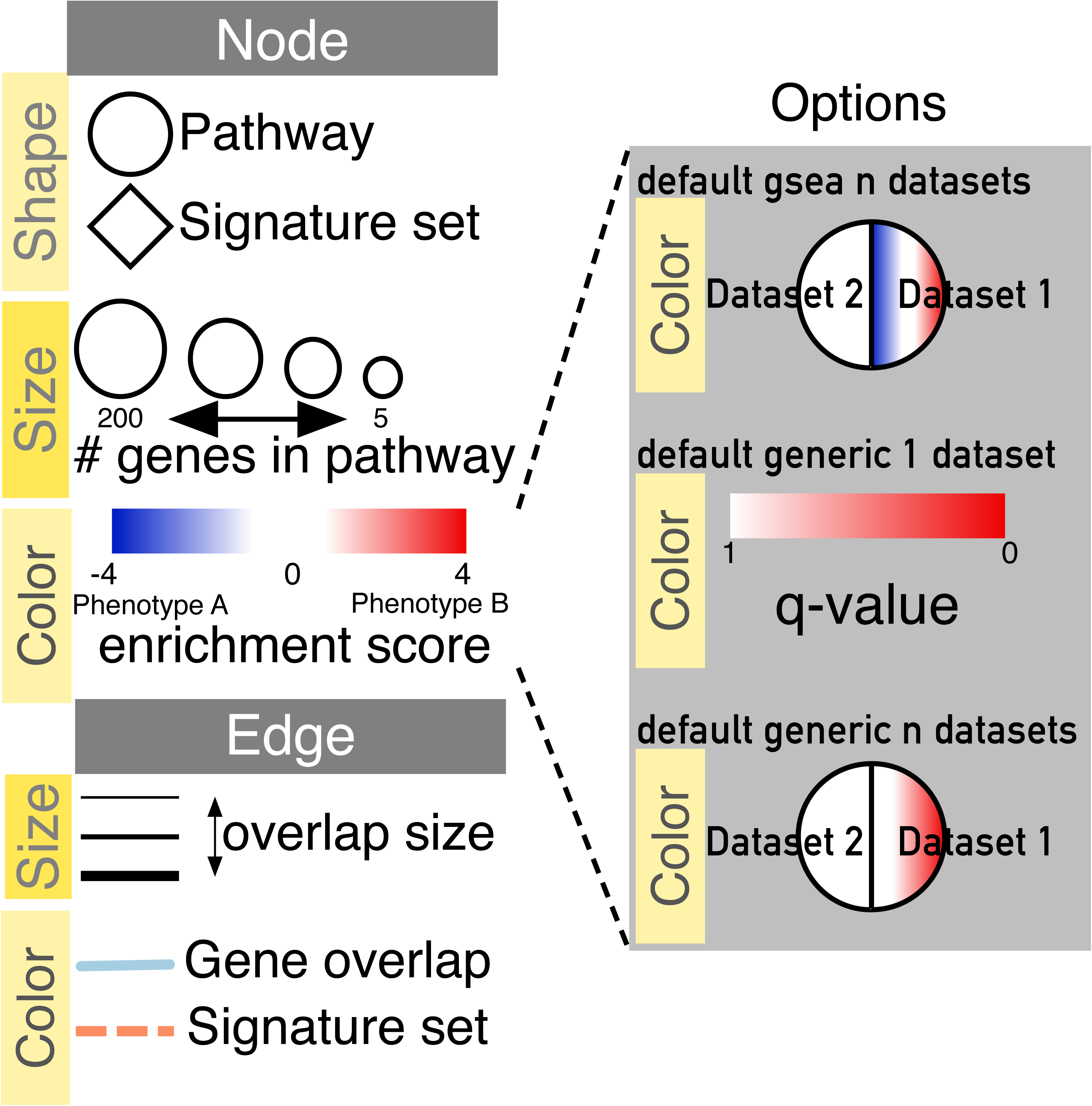

40 Save Cytoscape session (**TT37**). File**→**Save as. Navigate to the directory you wish to save the session and specify the desired file name.

#### TIMING

∼4 hours to analyze and annotate Enrichment Map using Cytoscape on Windows7 with 8GB of RAM and Java 8 (**TT38**).

## TROUBLESHOOTING

**TT1.** On launching GSEA on macOS for the first time, you may get the error “‘gsea.jnlp’ can’t be opened because it is from an unidentified developer”. Click on “Ok”. Instead of double clicking on the gsea.jnlp icon/file, right click and select “open”. The same error “’gsea.jnlp’ can’t be opened because it is from an unidentified developer” will appear but this time it will give you the option to “Open” or “Cancel”. Click on “Open”. After this initial opening, subsequent double clicks on gsea.jnlp will launch GSEA without any errors or warnings. If GSEA still fails to launch through the Java Web Start downloaded from the GSEA website, GSEA can be alternatively be launched from the command line. Go to the GSEA download site and download javaGSEA JAR file (the second option on the download site). Open a command line terminal. On macOS, the terminal can be found in Applications -> Utilities -> Terminal. On Windows type “cmd” in the windows program files search bar. Then navigate to the directory where the file javaGSEA.jar was downloaded using the command cd. For example, on macOS run “cd ∼/Downloads” if you downloaded the GSEA jar to your downloads folder. Run the command java -Xmx4G - jar gsea-3.o.jar where -Xmx specifies how much memory is given to GSEA.

**TT2.** It may take 5-10 seconds for GSEA to load input files. The files are loaded successfully once a message appears on the screen “Files loaded successfully: 2/2. There were no errors”.

**TT3.** GSEA also supplies its own gene set files that are accessible directly through the GSEA interface from the MSigDB resource^47,48^. These files do not need to be imported into GSEA. When you define the GMT file, the MSigDB gene set files can be found in the first tab “Gene Matrix (from website)” of the “Select one or more genesets” dialog. The latest versions of the MSigDB gene set files are in bold but previous versions can also be accessed. To select multiple gene set files, use multi-file select by simultaneously clicking on the desired files and holding the control key on Windows or command on macOS.

**TT4.** The higher the number of permutations the longer the analysis will take. To calculate the FDR *q*-value for each gene set, the data is randomized by permuting the genes in each gene set and recalculating the p-values for the randomized set. This parameter specifies how many times this randomization is done. The more randomizations are performed, the more precise the FDR q-value estimation will be (to a limit, as eventually the FDR q-value will stabilize at the actual value). On a Windows machine with 16G of RAM and i7 3.4 GHz processor, an analysis with 10,100, 500, or 1000 randomizations on our example set with above defined parameters takes 155, 224, 544, and 1012 seconds, respectively.

**TT5.** GSEA has no progress bar to indicate estimated time to completion. A run can take a few minutes or hours depending on your data size and computer speed. Click on the “+” in the bottom left corner of the screen to see messages such as “shuffleGeneSet for GeneSet 4661/4715 nperm: 1000” (circled in red at the bottom of Figure 2). This message indicates that GSEA is shuffling 4,715 gene sets 1,000 times each, 4,661 of which are complete. Once the permutations are complete, GSEA generates the report.

**TT6.** The error message “Java Heap space” indicates that the software has run out of memory. Another version of GSEA is needed in case you are running the Web Start application. There are multiple options available for download from the GSEA website. You can download a webstart application that launches GSEA with 1, 2, 4, or 8GB. Upgrade to a webstart that launches with more memory. If you are already using the webstart that launches with 8GB then you require GSEA JAVA jar file which can be executed from the command line with increased memory (see TT1 for details).

**TT7.** If the GSEA software is closed, you can still see the results by opening the working folder and opening the ‘index.html’ file. Alternatively, you can re-launch GSEA, and click on “Analysis history”, then “History” and then navigate to date of your analysis. Although all analyses, regardless of where the results files were saved, are listed under history, it is organized by date the analysis was run. If you can’t remember when you ran a specific analysis, then you may have to manually search through a few directories to find the desired analysis.

**TT8.** When running GSEA with expression data as input (instead of a pre-calculated rank file), a phenotype label (i.e. biological condition or sample class) is provided as input for each sample and specified in the GSEA ‘cls’ file. When running GSEA, the two phenotypes to compare for differential gene expression analysis are specified and these phenotypes are used in the pathway enrichment result files. In contrast, in a GSEA preranked analysis (i.e. when a ranked gene list is provided by the user), GSEA automatically labels one phenotype “na_pos” (corresponding to enrichment in the genes at the top of the ranked list, where ‘na’ means the phenotype label ‘not available’) and the other “na_neg” (corresponding to enrichment in the genes at the bottom of the ranked list). This convention is also used by the Enrichment Map software, designating the first phenotype as “positive” and the second phenotype as “negative”.

**TT9.** Check the number of gene sets that were analyzed. If the number is low (e.g. low hundreds), it could indicate gene ID mapping problems.

**TT10.** To simplify loading g:Profiler results into Enrichment Map and populating the correct fields in the EM interface, place the g:Profiler results file and gene set file (i.e. Supplementary_Table4_gprofiler_results.txt and Supplementary_Table5_hsapiens.pathways.NAME.gmt) into a directory together.

**TT11.** Although an individual g:Profiler analysis only has one phenotype, it is possible to modify a single results file to contain two analyses. This is relevant when the phenotypes are mutually exclusive. For the analysis you want to associate with the additional phenotype (which would correspond to down-regulated genes in GSEA PreRanked, thus encoded as “negative”) open the g:Profiler results file (preferably in a spreadsheet so you can easily modify a single column). The fifth column specifies the phenotype. Update the column to have the value of “-1” for each result in the file. Open the second analysis file. Copy all the results from the second file and paste them into the updated negative g:Profiler file. Save the file and use it as the g:Profiler enrichment results file in the EM interface instead of the original results files. Pathways corresponding to two phenotypes will be colored red and blue in the resulting enrichment map. One limitation with this approach is that a pathway cannot be included in both positive and negative sets.

**TT12.** Moving the slider to the left (or right) will adjust the underlying similarity statistic threshold to make the resulting network sparser (denser). The slider is set with predefined defaults but users can fine-tune the similarity metric by selecting ‘Show advanced options’ at the bottom of the ‘Create Enrichment Map’ panel. Predefined values appear as tick marks on the slider and include Jaccard > 0.35, Jaccard > 0.25, combined > 0.375, overlap > 0.5, overlap > 0.25.

**TT13.** If you specify a directory that contains multiple GSEA results rather than an individual GSEA results folder, EM will treat every GSEA results folder as its own data set. This enables easy multi-data set analyses. If you only wanted one data set but inadvertently selected the directory containing multiple GSEA results instead of selecting an individual folder, simply select the data sets you do not want to use and click on the trash can at the top of the EM input panel.

**TT14.** Every GSEA analysis generates a random number that is appended to the names of the files and directories. The number will be different for every new analysis.

**TT15.** If Enrichment Map cannot find the original GMT file used in the GSEA analysis, it will use a filtered GMT file found in the GSEA ‘edb’ results directory. Enrichment map will not be able to find your original GMT file if you have moved it since running GSEA analysis. Although it is a GMT file, it has been filtered to contain only genes found in the expression file. If you use this filtered file, you will get different pathway connectivity depending on the expression data being used. We recommend using original GMT file used for the GSEA analysis and not the filtered one in the results directory.

**TT16.** To annotate the phenotypes in the Enrichment Map heat map, the specified phenotype labels need to exactly match the GSEA CLS file.

**TT17.** If you load the CLS file prior to specifying the phenotypes, EM will automatically guess the phenotypes from the class file. If your class file specifies more than two phenotypes, EM will choose the first two phenotypes defined in the file.

**TT18.** To set the threshold to a small number, select ‘Scientific Notation’ and set a q-value cutoff such as 1E-04.

**TT19.** Multiple genes separated by spaces can be entered into the search bar. Any pathway that contains the gene will be selected and highlighted in the network. Adding keywords “AND” into the search bar will show only pathways that contain all genes in the search query. If the analysis was not done using gene symbols then you will not be able to search by gene symbols. Instead use the identifier the analysis was based on, for example Entrez gene ID or Ensembl gene ID.

**TT20.** If there are very few records in the node table make sure that no nodes are selected in the network. Click on the gear icon and change the setting from “Auto” to “Show all”.

**TT21**. If no expression file is given to Enrichment Map it will automatically create a dummy expression file where any gene found in the enrichment file will be given a placeholder expression value of 0.25, and any gene found in a pathway but not found in the enrichment results file be assigned a placeholder expression value of NA. Therefore clicking on any node in the enrichment map will show the genes used for the analysis as well as genes in the pathway but not part of the query set.

**TT22.** The leading edge can be displayed only if the rank file is provided when the network is built. The rank file supplied needs to be identical to the one used for the GSEA analysis for the leading edge calculation to be accurate.

**TT23.** In case of multiple conditions or conditions with variable expression profiles (e.g. cancer patient samples), hierarchical clustering tends to generate a more informative visualization.

**TT24.** If the heat map columns are not colored for a GSEA analysis, make sure the phenotype names specified in the Enrichment Map input panel match the class names specified in the class file (MesenchymalvsImmunoreactive_RNA-Seq_classes.cls)

**TT25.** Leading edge is only available for GSEA analyses. The option will only appear if the Enrichment map was built with GSEA results and a rank file was specified.

**TT26.** There are many different layout algorithms available in Cytoscape that can be used for Enrichment Map. We recommend using an edge weighted layout, which considers the overlap score between pathways. Most layouts offer the ability to organize just the selected nodes (except for yFiles layouts). Experiment with different layouts to see which works best with your data. If you do not like layout results simply press command-Z on macOS or ctrl-Z on Windows or click on Edit --> Undo to revert to the previous view.

**TT27.** If particular non-informative words keep appearing in the labels generated by AutoAnnotate, try adjusting the WordCloud normalization factor. The significance of each word is calculated based on the number of occurrences in the given cluster of pathways. This causes frequent words such as “pathway” or “regulation” to be prominent. By increasing the normalization factor, we reduce the priority of such recurrent words in cluster labels. If that doesn’t help you can add the non-informative words to the WordCloud word exclusion list.

**TT28.** If a specific character is used to separate words besides space (for example “-“ or “|”), it should be added as a delimiter in the WordCloud app. Launch the WordCloud App (Apps**→**WordCloud). In the WordCloud input panel expand Advanced options. Click on “Delimiters.”. Add your delimiters. Click on OK. In the AutoAnnotate input panel click on the menu button. Select “Recalculate Labels.” for this change to take effect.

**TT29.** The default parameters are likely to work well with Enrichment Map, however there are many parameters within the AutoAnnotate app that can fine-tune the results. See the AutoAnnotate user manual at http://www.baderlab.org/Software/AutoAnnotate/UserManual.

**TT30.** The number of nodes in a cluster determines label size by default. Thus, the cluster size may related to pathway popularity instead of importance in the experiment. Annotation labels can all be set to the same size by unchecking the option “Scale font by cluster size” in the AutoAnnotate results panel.

**TT31.** Once you click on “Collapse All” a pop-up window will show the message “Before collapsing clusters please go to the menu Edit->Preferences->Group preferences and select “Enable attribute aggregation”. There is no need to adjust this parameter repeatedly. Click on “Don’t ask me again” and “OK” if you have set this parameter previously.

**TT32.** For large networks, collapsing and expanding may take time. For a quick view of the collapsed network you can create a summary network by selecting the “Create summary Network…”. There are two options for the summary network: “clusters only” which creates a summary network with just the circled clusters, or “clusters and unclustered nodes” which creates a summary network that also includes the singleton nodes not part of any cluster.

**TT33.** If the nodes in the resulting collapsed network are grey then you forgot to enable attribute aggregation. Expand clusters and refer to **TT31** before collapsing again.

**TT34.** In image export, only the visible part of the map will be exported. Make sure that the entire network is visible on your screen before exporting.

**TT35.** Vector-based PDF and SVG formats are recommended for publication quality figures because they can zoom without losing quality. Either file type can be edited using software packages such as Adobe Illustrator or Inkscape. The PNG file format is recommended for high-quality online images while the JPG format is not recommended because it may lead to visual artefacts due to compression.

**TT36.** The creation parameters panel only shows the parameters that were used at network creation. If you modified the network using filters or the EM slider bars you will have to update the changed thresholds accordingly.

**TT37.** If a session is saved that contains a collapsed Enrichment Map, it will automatically be expanded before it is saved. Depending on the size of the network this might take a few minutes. Enrichment Map will not automatically collapse nodes that were previously collapsed when reopening the session. To keep them collapsed, recollapse them using the AutoAnnotate app as done previously.

**TT38.** The time required for this protocol depends on the amount of manual curation and visual organization spent on the enrichment map. Smaller networks are generally easier to organize. Before spending the time laying out the final network, it is worth revisiting the enrichment analysis, exploring the network fully and selecting the parts to emphasize in a final publication quality figure.

## Author contributions

J.R., R.I., V.V., A.R., D.M., G.D.B. wrote the manuscript. R.I created the step-by-step protocols, figures, R scripts and R notebooks, except for g:Profiler (J.R.). M.K, C.T.L. developed Enrichment Map 3.0, and AutoAnnotate Cytoscape apps. L.W., M.M., J.W., C.X., V.V. tested the protocol. All authors read and approved the final manuscript.

## Acknowledgements

The authors are grateful to Jill Mesirov for comments on the manuscript. This project was supported by an Investigator Award to JR from the Ontario Institute for Cancer Research through funding provided by the Government of Ontario. This work was supported by U.S. National Institutes of Health grants P41 GM103504, R01 GM070743, U41 HG006623, R01 CA121941 to GDB.

### BOX 1 - Definitions

**Pathway** – genes that work together to carry out a biological process.

**Gene set** – A set of related genes. A ‘pathway gene set’ includes all genes in a pathway. Gene sets can be based on various relationships between genes, such as cellular localization (e.g., nuclear genes) or enzymatic function (e.g., protein kinases). Details such as protein interactions are not included.

**Gene list of interest** – the list of genes derived from an –*omics* experiment that is input to pathway enrichment analysis.

**Ranked gene list** – In many –*omics* data (e.g. RNA-seq for gene expression) genes can be ranked according to some score (e.g. level of differential expression) to provide more information for pathway enrichment analysis. Pathways enriched in genes clustered at the top of a ranked list would score higher than if the pathway genes are randomly scattered across the ranked list.

**Pathway enrichment analysis** – a statistical technique to identify pathways that are significantly represented in a gene list or ranked gene list of interest.

**Multiple testing correction** – Thousands of pathways may be individually tested for enrichment and this could lead to significant enrichment p-values appearing by chance alone. Multiple testing correction is a statistical technique to correct the p-values from individual enrichment tests to address this problem and reduce the chance of false positive enrichment (**Box 3**).

### BOX 2 - Experimental design and data quality

Pathway enrichment analysis benefits greatly from careful experimental design. Otherwise the analysis may reveal apparently meaningful results caused by experimental biases or other confounders.

**Experimental conditions.** The experimental conditions must be well defined such that the major variations observed are responses which the experimenter would like to monitor, related to the biological question of interest *(e.g.,* tumour *vs.* normal, treated *vs.* untreated, comparison of four disease subtypes, time series).

**Number of replicates.** Biological replicates are independently processed samples obtained from distinct organisms or cell lines that are required for measuring variability across samples and to compute statistics *(e.g.,* p-values). Lack of replication (i.e. one sample per group) will not permit robust estimation of the significance of the signal. Insufficient replication may result in lack of signal in the data (e.g. no significant differentially expressed genes). The larger the variation in the set of samples, the more biological replicates are needed to accurately measure signal. For systems with lower variability (i.e. model organisms with the same genetic background in controlled laboratory conditions, or stable cell lines derived from the same clone), at least 3-4 biological replicates are recommended per condition for differential analysis with variance stabilization normalization. Variance stabilization uses a global statistical model to “stabilize” gene-wise variance estimates to reduce inaccuracies resulting from few replicates. For experiments with higher variability (e.g. tumor samples), more replicates are required; ideally, a pilot experiment followed by formal power calculations should be used to determine the minimal number of replicates required to identify signal of differentially expressed genes or enriched pathways. Technical replicates comprising repeated experiments of the same samples are usually not needed for well-established experimental techniques that have low technical variability, like RNA-seq, but can be helpful for novel techniques.

**Confounding factors.** Differences in factors not related to the experimental question should be avoided or at least balanced across conditions so that statistical techniques such as generalized linear models can correct for each factor. Common factors include sequencing batch, nucleic acid extraction protocol, subject age and many others. Otherwise it may be impossible to accurately separate the experimental signals coming from the experimental response and confounding factors. Knowing important factors in advance supports correct experimental design. Statistical exploratory analyses such as clustering or principal component analysis (PCA) can help identify unknown factors. For example, cases and controls are expected to cluster separately and not by processing batch.

**Outliers.** Outlier samples may considerably differ from others because of major experimental or technical problems, such as contamination or sample mix-up. Alternatively, they may present extreme biological features, such as tumour samples with exceptionally aggressive phenotypes. Unbiased identification of outlier samples is possible using statistical techniques such as PCA or clustering. Pathway enrichment analysis should be performed with and without outliers to ensure robust results. Systematic removal of outliers may be justified to reduce variability in the experiment.

**Experimental sensitivity.** Some experimental methods can be tuned to be more or less sensitive. For instance, the number of reads in RNA-seq experiments influences downstream analysis. For quantifying gene expression in a biological system with modest variability and testing differential expression with variance stabilization, at least three to five replicates and ten million mapped reads are required^49^. Substantially greater sequencing depth, such as 50-100 million mapped reads, is required to investigate splice isoforms, to detect low-expressed genes or for samples with complex cellular mixtures such as surgical resection specimens.

### BOX 3 - Statistical Tests used in Pathway Enrichment Analysis

A common statistical test used for pathway enrichment analysis of a gene list is a Fisher’s exact test, based on the hypergeometric distribution. It determines whether the fraction of genes of interest in the pathway is higher compared to the fraction of genes outside the pathway (i.e. background set). Since this test was first introduced^50^, many improved tests have been developed^51^ that take advantage of continuous experimental scores and avoid applying arbitrary thresholds. We categorize types of statistical enrichment tests as follows:

1. **Ranked vs. non-ranked tests.** Ranked tests take as input a ranked gene list, while non-ranked tests such as Fisher’s exact test take as input a gene list of interest. Ranked tests are preferable for experiments that produce meaningful ranks such as differential gene expression, because arbitrary thresholds can be avoided. Non-ranked tests are preferable for experiments that naturally generate a gene list of interest (e.g., somatic mutations in cancer, proteins that interact with a bait protein). Examples of ranked tests include the modified Fisher’s exact test implemented in the g:Profiler ‘ordered query’ option, and the modified Kolmogorov-Smirnov test implemented in GSEA.

2. **Exact vs. permutation-based tests.** Exact tests employ a mathematical model (e.g. a distribution) to directly compute an exact p-value. Permutation-based tests utilize data resampling to estimate an empirical p-value, typically expressed as number of permutations with results as good as or better than the ones observed for real data, divided by the number of permutations. For example, in a case-control study, we can randomize the case and control labels 1,000 times, each time repeating the pathway enrichment analysis to see how frequently we observe an equal or stronger pathway enrichment signal. Permutation tests can be customized to consider specific data properties and biases. Exact tests, if applicable, are preferable as these can quickly compute accurate p-values.

3. **Competitive vs. self-contained tests.** Competitive tests determine whether the gene list of interest is enriched in pathways relative to all genes in the background set. Thus, each pathway “competes” for enrichment in the gene list against genes of the background set. In contrast, self-contained tests calculate statistics uniquely at the pathway level, ignoring genes of the background set. For instance, a self-contained test can evaluate whether the gene expression within a given pathway is different in case samples compared to control samples^51^. Competitive pathway enrichment analysis is most popular and is usually appropriate for gene expression data. However, self-contained tests must be used if single gene differences are not significant and need to be pooled at the pathway gene set level to identify signal, for example when analysing rare gene mutations or other data with low per-gene counts^52^. Hybrid approaches may be preferable to self-contained tests in specific circumstances. For instance, for rare copy number variation (CNV) data, correcting a self-contained test for global CNV burden leads to more specific biological results^42^. Finally, competitive enrichment tests such as Fisher’s exact test ignore correlation among genes while modified competitive tests such as Camera^53^ consider these and thus typically produce more rigorous results. Self-contained tests do not present this issue.

In summary, if genes in your data can be ranked, a ranked test should be used. Fisher’s exact test is generally chosen for non-ranked gene lists. A competitive test is adequate in most cases, unless the signal at the gene-level is weak.

**Multiple test correction.** Repeated statistical testing used in a typical pathway enrichment analysis will result in some apparently significant p-values by chance alone. To correct this, multiple testing correction methods systematically reduce the significance of each p-value derived from a series of tests. The most commonly used method is the Benjamini-Hochberg False Discovery Rate (BH-FDR, or often simply FDR)^20^. It is based on a step-down procedure that estimates the fraction of falsely enriched pathways over enriched pathways, using the uncorrected p-value threshold and the number of tests. For instance, given that 100 pathways have enrichment p-value < 0.05 having a FDR of 5% at p-value < 0.05 means that five of those pathways are expected to be falsely enriched. As an alternative, the classical Bonferroni multiple testing correction adjusts the significance threshold by dividing it by the number of tests. Practically, the method multiplies each uncorrected p-value by the number of conducted tests and applies a significance cut-off *(e.g.,* a p-value of 0.001 will become an insignificant q-value 0.1 if 100 pathways have been tested). This technique ensures that the probability of selecting *at least one* falsely enriched pathway is below the corrected p-value threshold. Bonferroni correction is typically considered overly conservative for differential gene expression and pathway enrichment analysis because some fraction of false positive findings can be tolerated. Importantly, both Bonferroni and BH-FDR assume tests are independent, while pathways are typically not independent because of overlapping genes and cross-talk. Therefore, BH-FDR estimates for pathway analysis can be inaccurate, although practically they are still useful for filtering and hypothesis generation and thus are routinely used.

### BOX 4 - Pathway Enrichment Analysis Resources

**Pathway databases**

We describe a selection of large, open-access and conveniently accessible pathway databases that offer the maximal value for pathway enrichment analysis. Hundreds of pathway databases are available for many purposes^54^.

**Databases of gene sets**

- Gene Ontology (GO)^27^ - GO provides a hierarchically organized set of thousands of standardized terms for biological processes, molecular functions and cellular components as well as curated and predicted gene annotations based on these terms for multiple species. Biological process annotations are the most commonly used resource for pathway enrichment analysis.
- Molecular Signatures Database (MSigDB)^47,48^ -MSigDB is a collection of gene sets based on GO, pathways, curation, individual –*omics* studies, sequence motifs, chromosomal position, oncogenic and immunological expression signatures, and various computational analyses. Aggregate ‘hallmark’ gene sets are available as a relatively non-redundant collection (http://www.msigdb.org). It is created by the team that makes GSEA, but can be used with any pathway enrichment method.

**Detailed biochemical pathway databases.** These databases are maintained by human curators who manually collect detailed pathway information, including biochemical reactions, gene regulatory events and other gene interactions. The information can be exported or converted to gene set format.

- Reactome^28^ - the most actively updated public database of human pathways (http://www.reactome.org)
- Panther^21^ - human signalling pathways (http://pantherdb.org/pathway)
- NetPath^30^ - human signaling pathways with a focus on cancer and immunology (http://www.netpath.org/)
- HumanCyc^29^ - human metabolic pathways (http://humancyc.org/)
- NCI PID - human cancer related signaling pathways. No longer updated.
- KEGG^55^ - most useful for its intuitive pathway diagrams. Contains multiple types of pathways, some of which are not normal pathways, but are rather disease associated gene sets, such as “pathways in cancer” (http://www.genome.jp/kegg/)

**Pathway meta-databases.** These databases collect detailed pathway descriptions from multiple originating pathway databases.
- Pathway Commons^35^ - collects information from other pathway databases and provides it in a standardized format (http://www.pathwaycommons.org).
- WikiPathways^37^ - a community-driven collection of pathways that also includes pathways exported from other databases (http://www.wikipathways.org/). **Pathway Enrichment Analysis Tools** Hundreds of pathway enrichment analysis tools exist, although many of them rely on out-of-date pathway databases or do not present any unique feature compared to the most commonly used tools. The following are free pathway enrichment analysis software tools that we recommend based on their ease of use or unique features:

- g:Profiler^4,26^ – analyzes gene lists using Fisher’s exact test and ordered gene lists using a modified Fisher’s test. It provides a graphical web interface and access via R and python programming languages. The software is frequently updated and the gene set database can be downloaded as a GMT file (http://biit.cs.ut.ee/gprofiler).
- Gene Set Enrichment Analysis (GSEA)^5,56^ – analyzes ranked gene lists using a permutation-based test. The software runs as a desktop application (http://software.broadinstitute.org/gsea).
- Genomic Regions Enrichment of Annotations Tool (GREAT)^41^ – analyzes genomic regions, such as DNA binding sites, and links them to nearby genes (http://bejerano.stanford.edu/great/public/html/)
- Camera^53^ - analyzes gene lists and corrects for inter-gene correlations such as gene co-expression; available as part of the limma package in Bioconductor (https://bioconductor.org/packages/release/bioc/html/limma.html).
- GOseq^57^ (Advanced tool, requires programming) – this R Bioconductor package analyzes gene lists from RNA-seq experiments by correcting for user-selected covariates such as gene length (https://bioconductor.org/packages/release/bioc/html/goseq.html).

**Visualisation tools**

- Enrichment Map^7^ – this Cytoscape^6^ app visualizes the results from pathway enrichment analysis, eases interpretation by displaying pathways as a network where overlapping pathways are clustered together to identify major biological themes in the results (http://apps.cytoscape.org/apps/enrichmentmap).
- ClueGO^34^ – This Cytoscape app is conceptually similar to Enrichment Map. It includes a GO-based pathway enrichment analysis feature using Fisher’s exact test.
- PathVisio^38^ – this desktop application displays genomic data on a pathway diagram (https://www.pathvisio.org).

### BOX 5 - Topology-Aware Pathway Enrichment Analysis Methods

Most pathway enrichment analysis methods treat all genes in a pathway uniformly and ignore gene interactions. In contrast, topology-aware methods explicitly model the interactions between genes. CePa^58^, GANPA^59^, and THINK-Back^60^ use physical gene interactions or co-expression networks to assign a weight to each gene in each pathway. Weights can be derived from measures of the gene importance in the network such as degree, the number of gene connections, and betweenness centrality, and can be integrated into a traditional pathway enrichment analysis method such as GSEA. Methods like SPIA^61^, Pathway-Express^62^, and EnrichNet^63^ generate an enrichment score for the entire pathway that considers pathway regulatory interactions, such as activation and inhibition. While useful and potentially more accurate, regulatory gene interactions are available for fewer genes compared to physical interactions networks and co-expression.

## Supplementary Materials

1. Supplementary_Table_1_Cancer_drivers.txt

2. Supplementary_Table2_MesenvsImmuno_RNASeq_ranks.rnk

3. Supplementary_Table3_Human_GOBP_AllPathways_no_GO_iea_July_01_2017_symbolgmt

3. Supplementary_Table4_gprofiler_results.txt

4. Supplementary_Table5_hsapiens.pathways.NAME.gmt

5. Supplementary_Table6_TCGA_OV_RNAseq_expression.txt

6. Supplementary_Table7_gsea_report_for_na_pos.xls

7. Supplementary_Table8_gsea_report_for_na_neg.xls

8. Supplementary_Table9_TCGA_OV_RNAseq_classes.cls Files for supplementary protocols:

9. Supplementary_Table10_TCGA_Microarray_rmanormalized.txt

10. Supplementary_Table11_Microarray_classdefinitions.txt

11. Supplementary_T able12_TCGA_RNASeq_rawcounts.txt

12. Supplementary_Table13_RNASeq_classdefinitions.txt

## Supplementary Protocols

This protocol processes RNA-seq data using the R programming environment and specialized packages from Bioconductor to create genes lists. The scripts are available for download and novice users can copy and paste commands into the R console. To create gene expression data for **Protocol 1B**, we downloaded gene expression data from the Ovarian Serous Cystadenocarcinoma project of The Cancer Genome Atlas (TCGA)^64^, http://cancergenome.nih.gov via the Genomic Data Commons (GDC) portal^65^ on 201706-14 using TCGABiolinks R package^66^. The data includes 544 samples available as RMA-normalized microarray data (Affymetrix HG-U133A), and 309 samples available as RNA-seq data, with reads mapped to a reference genome using MapSplice^67^ and read counts per transcript determined using the RSEM method^68^. RNA-seq data are labeled as ‘RNA-Seq V2’, see details at: https://wiki.nci.nih.gov/display/TCGA/RNASeq+Version+2). The RNA-SeqV2 data consists of raw counts similar to regular RNA-seq but RSEM (RNA-Seq by Expectation Maximization) data can be used with the edgeR method.

### Equipment

#### Hardware requirements

- A recent personal computer with at least 8 gigabytes of memory (RAM).

#### Software requirements

- The R statistical computing environment (http://www.r-project.org/). We suggest using the integrated development environment RStudio (https://www.rstudio.com/).
- Required R packages: Biobase, Limma and GSA are available from Bioconductor (https://www.bioconductor.org/)

#### Data requirements

- Supplementary_Table10_TCGA_Microarray_rmanormalized.txt corresponds to the RMA normalized Affymetrix mRNA transcript expression data for serous ovarian cancer samples as downloaded from the GDC portal^65^. Normalization is required to compute differential gene expression values across subtypes, as performed in **Supplementary Protocol 1A**.
- Supplementary_Table12_TCGA_RNASeq_rawcounts.txt corresponds to read counts per mRNA transcript determined using the RSEM method. These counts can be used to compare gene expression between subtypes using the edgeR analysis tool, as performed in **Supplementary Protocol 1B**. The counts are not pre-normalized and the normalization step using edgeR is part of the protocol.
- Supplementary_Table11_Microarray_classdefinitions.txt and Supplementary_Table13_RNASeq_classdefinitions.txt define the subtype classification of ovarian cancer samples (immunoreactive, mesenchymal, differentiated, proliferative) (Verhaak et al.^44^ supplementary table 1, column 3). This information is used to extract two subgroups of interest, mesenchymal and immunoreactive.

### Equipment Setup

- Download and install R from http://cran.r-project.org/
- Download and install RStudio from https://www.rstudio.com/ (optional, but recommended)
- Launch R or RStudio
- Install required Bioconductor packages. Enter the following commands in the R command line (also see https://www.bioconductor.org/install/):

- source("http://www.Bioconductor.org/biocLite.R")
- biocLite("BiocUpgrade")
- biocLite(c("Biobase","limma",”edgeR”,”GSA”,”locfit”))
- install.packages(c(“pheatmap”, “RColorBrewer”, “gProfileR”, “RJSONIO”, “httr”))
- If the required packages are already installed you may receive a prompt to update these. The prompt window will ask you about updating the packages:

- Update all/some/none? [a/s/n]
- Type ‘a’ without quotes and hit enter.
- Load libraries into the R session using the example below. Loading of libraries is required every time R is re-opened.

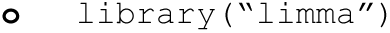
- All R files are available at https://github.com/BaderLab/EM-tutorials-docker/tree/master/Rscripts
- Alternately, R notebooks (use R markdown and creates a notebook similar to Jupyter notebooks) of this protocol are available at https://github.com/BaderLab/Cytoscape_workflows/tree/master/EnrichmentMapPipeline

### Data setup

- Download **Supplementary Tables 10**-**13** to a dedicated folder of your computer. The first step of your R script will change the working directory of R to this folder.
- As text editors sometimes add invisible characters to text copied from PDF files, copying and pasting from this document is not recommended. The R scripts or the R notebook available in the above URLs should be used instead.
- Setting the current directory and loading packages (libraries) are the required first and second steps of each protocol. These are needed each time a new session is opened in R.

### Supplementary Protocol 1 – create a gene list by analyzing gene expression data from RNA-seq using edgeR

This part of the supplementary protocol demonstrates filtering and scoring RNA-seq data using normalized RNA-seq count data with the edgeR R package. The protocol can be used to produce input data for pathway enrichment methods like g:Profiler, GSEA and others. This RNA-seq analysis protocol follows conceptually similar steps to microarray analysis shown above.

1. Load required Bioconductor packages into R.

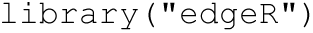

2. Load the expression data of 300 tumours, with 79 classified as Immunoreactive, 72 classified as Mesenchymal, 69 classified as Differentiated, and 80 classified as Proliferative samples. The TCGA counts data was retrieved from the Genomic Data Commons (GDC)^65^ database and contained counts per mRNA transcript determined using the RSEM method for 19947 transcripts and 300 samples.

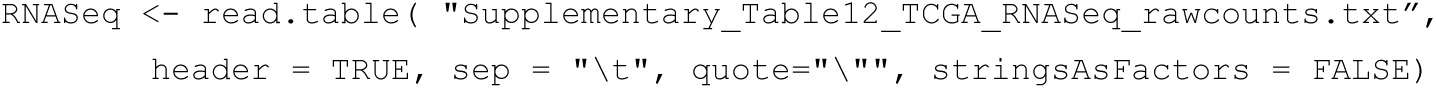

3. Load subtype classification of samples. To calculate differential expression, we need to define at least two sample classes. A common experimental design involves cases and controls but any two classes can be used. The current set of samples is divided into mesenchymal and immunoreactive classes (class definitions were obtained from Verhaak et al.^44^ Supplementary Table 1, third column). After loading the matrix, check that the column names of the expression matrix and class definitions are equal.

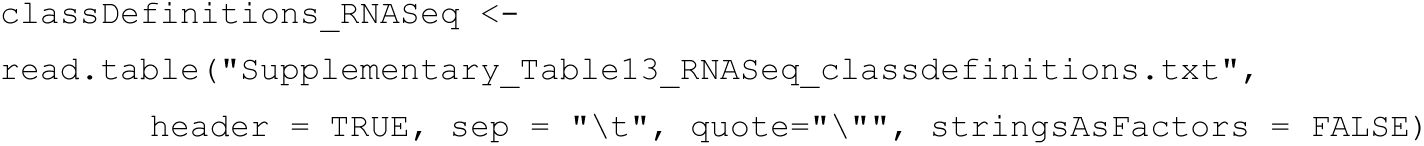

4. Filter RNA-seq reads. RNA-seq data are processed following the edgeR protocol^16^ that filters reads based on the counts per million (CPM) statistic. RNA-seq read counts are converted to CPM values and genes with CPM > 1 in at least 50 of the samples are retained for further study (50 is the minimal sample size in the classes). This step removes genes with very low read counts that are likely not expressed in the majority of samples and cause noise in the data. Note, CPM filtering is used to remove low counts while differential gene expression analysis is based on normalized read counts which are generated below (step 6).

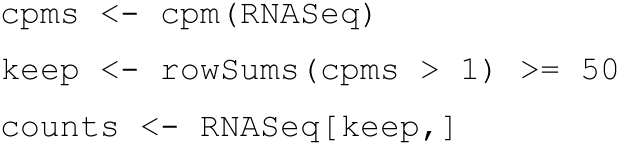

5. Data normalization, dispersion analysis is performed on the entire data.

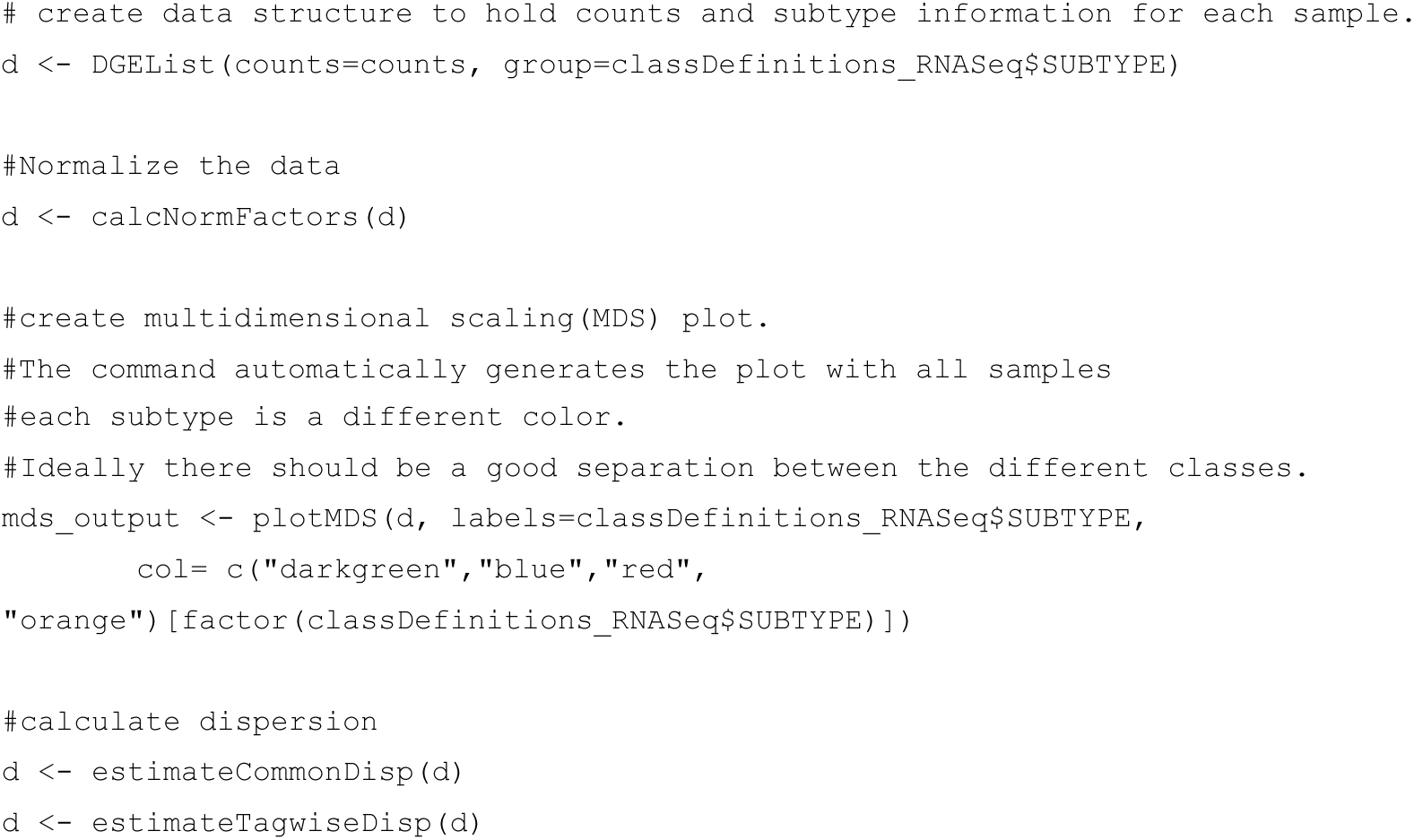

6. (Optional) Exclude genes with missing symbols or uncharacterized genes. In this example gene entries in the data containing ‘?’ or starting with LOC are excluded as they represent non-annotated genes or other loci that are not present in pathway databases. The frequency of these and other non-protein coding entries in the input data will depend on the database used to align the RNA-seq data.

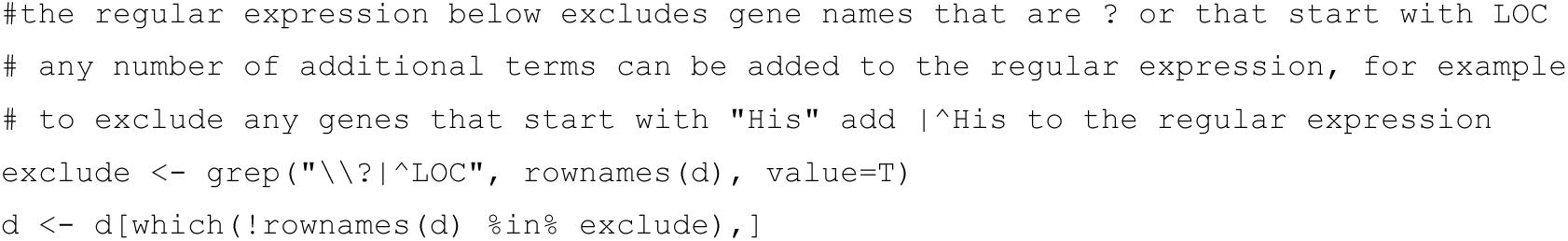

7. Differential expression analysis is performed with a simple design as described in the edgeR protocol^16^.

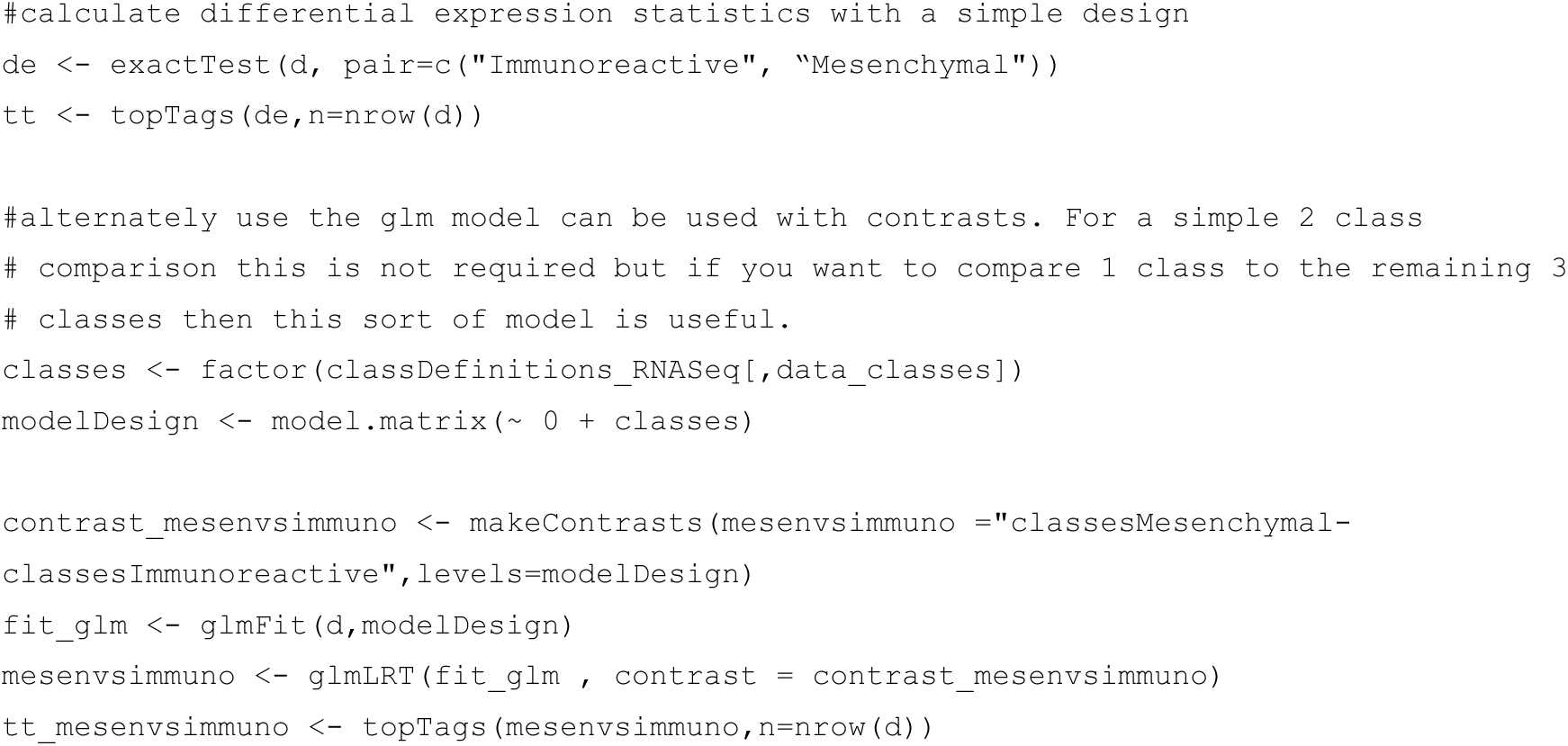

8a. Create the gene list for use in g:Profiler or another thresholded enrichment tool. The list may comprise all genes that have a significant *q*-value (code shown below), all significant and FDR-corrected up-regulated genes and all down-regulated genes separately, or some other combination of thresholds. Also see analogous step in the microarray protocol (Supplementary Protocol 2).

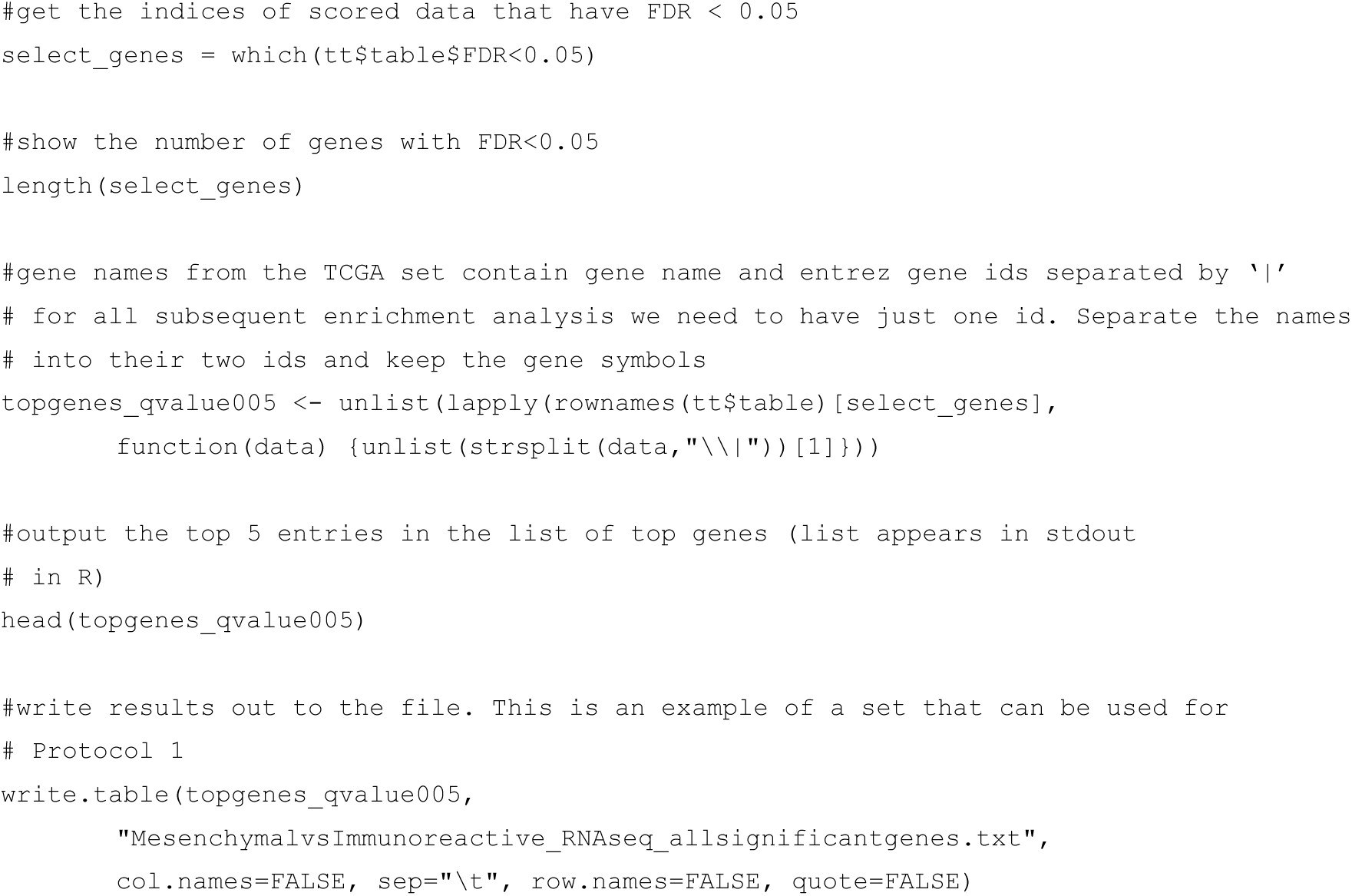

8b. Create a two-column rank (.RNK) file of all gene IDs and corresponding scores for input to GSEA pre-ranked analysis. One option is to rank genes by t-statistic of differential gene expression. GSEA will look for enrichment in the set of most differentially expressed genes at the top of the list as well as those at the bottom of the list. Genes at the top of the list are more highly expressed in class A of samples (e.g., mesenchymal) while genes at the bottom are highly expressed in class B (e.g., immunoreactive). An alternative score that we use here is computed by multiplying direction (sign) of fold change and logarithm of p-value for each gene.

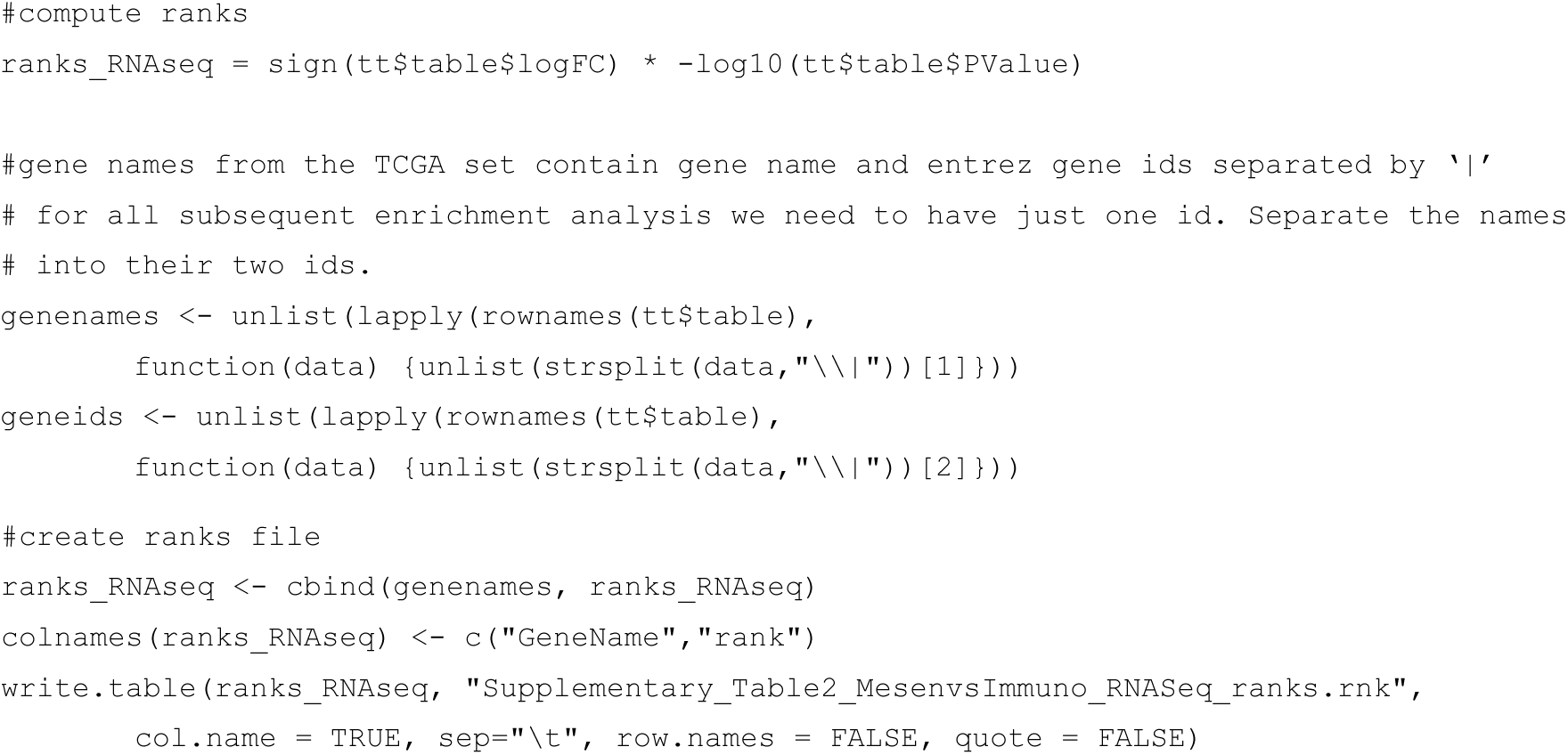

9a. Create an expression file for the enrichment map and save it to a file in the working folder. The optional expression file is similar to the expression matrix except for an additional column on the left edge of the matrix. The field often includes gene description however any text value can be added.

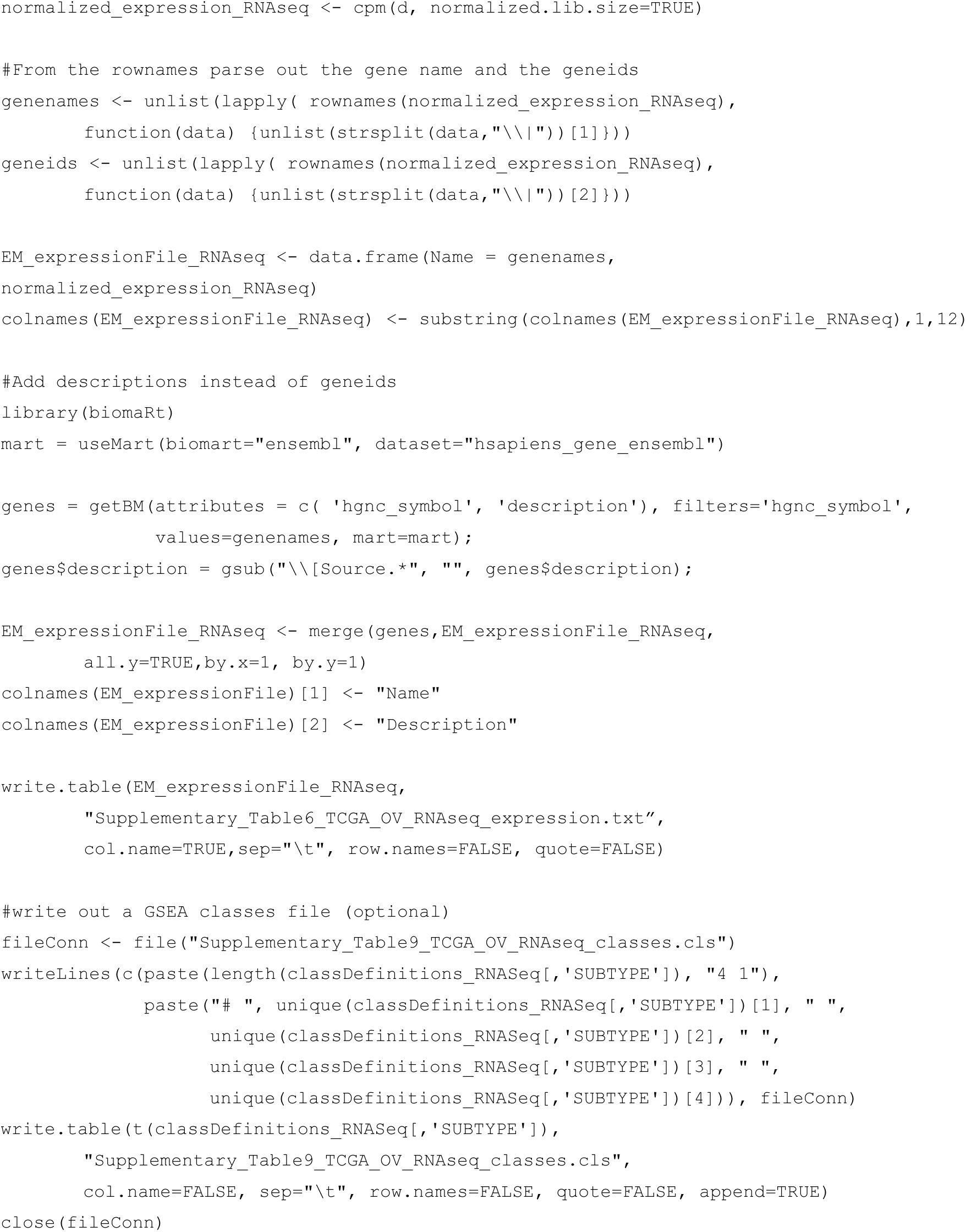

9b. Examine gene expression data using heat maps. Clustered heat maps can easily show the separation between sample classes, labeled by colors in the heat map header. By limiting to the most significantly differentially expressed list of genes (FDR *q*<0.05) we can verify whether the scoring accurately separates class A from class B.

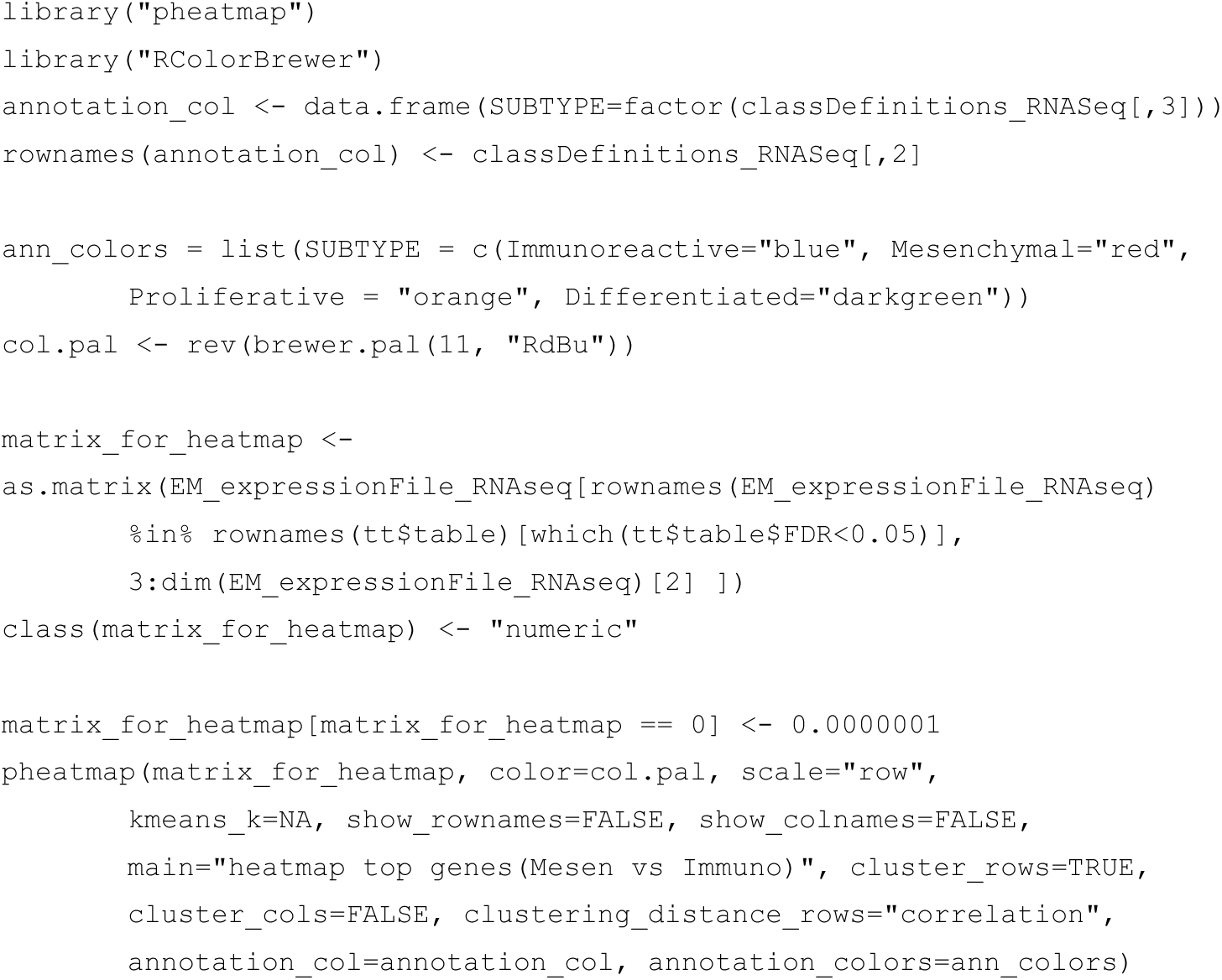

### Supplementary Protocol 2 – create a gene list by analyzing gene expression data from Affymetrix microarrays with Limma

This protocol demonstrates the generation of gene lists for pathway enrichment analysis using RMA-normalized gene expression data from Affymetrix microarrays for downstream pathway enrichment analysis with g:Profiler, GSEA and other similar tools. g:Profiler requires a ranked list of differentially expressed genes that are filtered according to a significance cut-off. GSEA requires a two-column tab-separated RNK file with a ranked list of all genes in the genome. In the RNK file, the first column specifies the gene name and the second column specifies a numeric score representing the level of differential expression. For both methods, the first step involves calculating a statistic for each gene that represents the difference in its expression levels between the two groups. This step is performed using the limma R package.

9. Load required Bioconductor packages into R

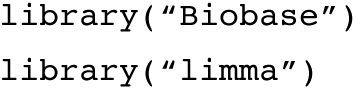

10. Set the working directory to the location of **Supplemental Tables 10-13**. The function getwd() shows the working directory and dir() shows its files.

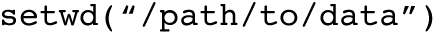

11. Load expression data into R. Minimally the expression set requires a gene name for each row and typically at least six expression values (three values in each compared class). Our data consists of 499 patients with 108 immunoreactive, 112 mesenchymal, 138 differentiated and 141 proliferative samples. After loading, use the command head(expressionMatrix) to verify that the matrix loaded correctly.

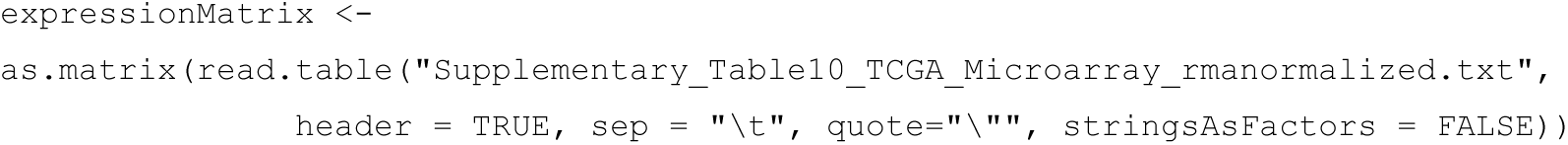

12. Load subtype classification of samples. To calculate differential expression, we need to define at least two classes of samples. A common experimental design involves cases and controls but any two classes may be used. The current data is divided into mesenchymal and immunoreactive classes (**Supplementary Table 11**, third column). After loading the matrix, check that the column names of the expression matrix and class definitions are equivalent.

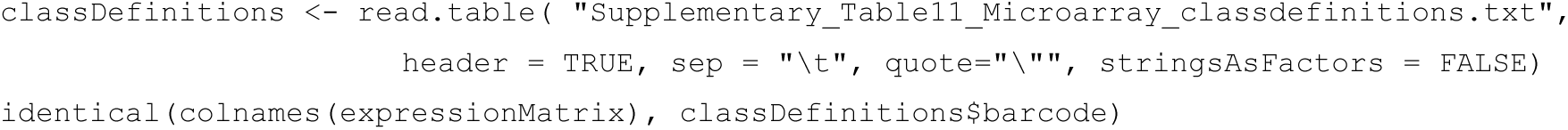

13. Format data and class definitions for limma. The expression data needs to be converted to an object of type ExpressionSet. The ExpressionSet must include a data matrix where rows are genes, columns are samples and each cell contains an expression value. Classes need to be defined as factors.

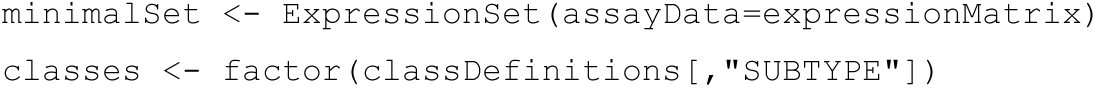

14. Create a model matrix with the defined classes.

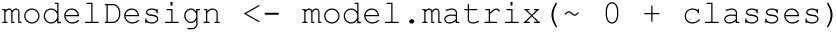

15. Fit the model to the expression matrix.

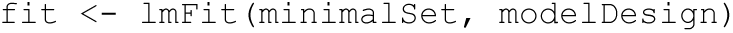

16. Create the contrast matrix. By specifying Mesenchymal first and Immunoreactive second, positive logFC and t-values refer to higher expression levels (up-regulation) in the Mesenchymal versus Immunoreactive samples

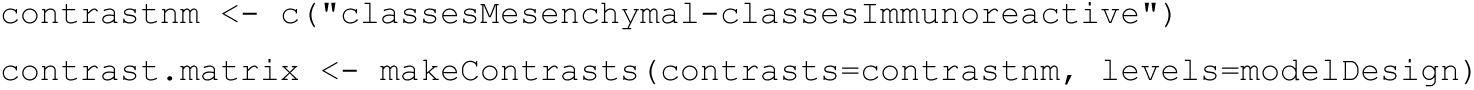

17. Model contrasts of gene expression. The following command models gene expression differences of each gene between the two groups of samples using linear regression and computes coefficients and standard errors.

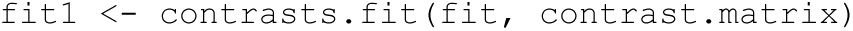

18. Compute differential expression statistics. Given a fitted linear regression model, the command generates a table containing the log fold change, average expression, t-statistic, *p*-value, adjusted *p*-value (*q*-value) and B statistic for each entity in the expression matrix using empirical Bayes statistics. The B-statistic represents the log-odds that the gene is differentially expressed, but it is based on a prior assumption of how many genes are differentially expressed in the data. Because of its reliance on this prior assumption, the adjusted p-value is preferentially used as an indicator of significant differential expression.

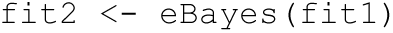

19. Generate a table with differentially expressed genes and adjust for multiple hypothesis testing using Benjamini-Hochberg False Discovery Rate. The table contains all genes ranked by p-value and shown with log fold change, average expression, t-statistic, p-value, adjusted p-value (*q*-value) and B-statistic.

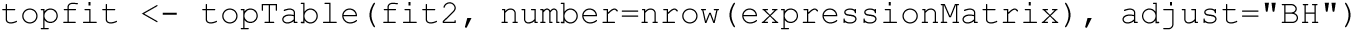

20a. Create the gene list for use in g:Profiler or another threshold-requiring enrichment tool. The list may comprise all genes that have a significant FDR *q*-value, all up-regulated genes with a significant FDR *q*-value, all down-regulated genes with a significant q-value, or some other combination of thresholds.

- To get all significant genes:

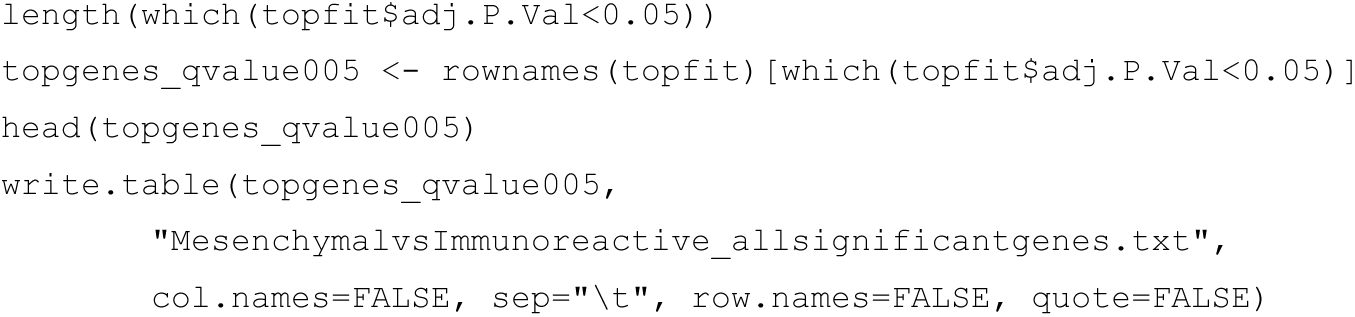

- Significantly up-regulated genes in mesenchymal samples have positive logFC and t-values.

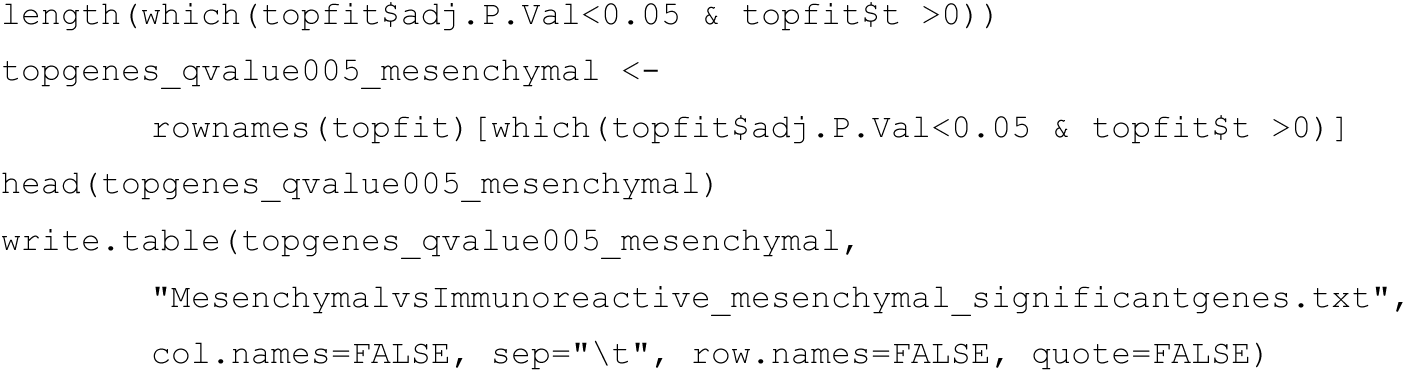

- Significantly up-regulated genes in immunoreactive samples have negative logFC and t-values.

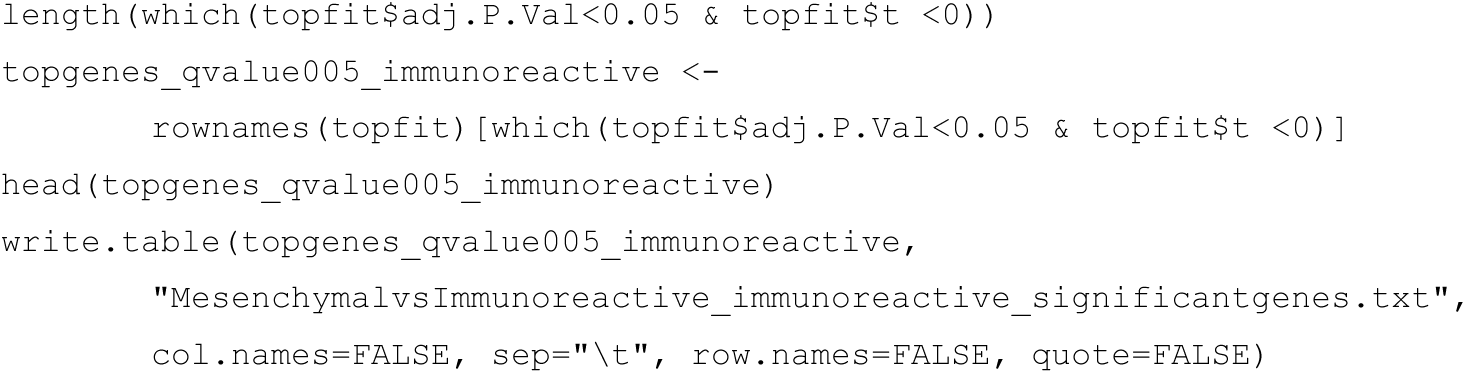

20b. Create a rank file for GSEA. To run GSEA in pre-ranked mode, you need a two column RNK file with gene/protein/probe name (column 1) and the associated score (column 2). The first column should contain the same type of gene IDs used in the pathway gene set (GMT) file. GSEA looks for enrichment in the top and bottom parts of the list by ranking the file using the t-statistic. The t-statistic indicates the strength of differential expression and is used in the p-value calculation. Other scores indicating the strength of differential expression may be used as well. GSEA ranks the most up-regulated genes at the top of the list and the most down-regulated at the bottom of the list. Genes at the top of the list are more highly expressed in class A compared to class B, while genes at the bottom of the list are higher in class B. In this workflow, a positive t-value means a higher gene expression in the Mesenchymal samples compared to the Immunoreactive samples (variable `constrastnm`). The following commands create a data frame with gene IDs and t-statistics, remove lines with missing gene IDs, and store the result as a RNK file. An additional step is usually required in analysis of Affymetrix microarray data as genes are represented with multiple probe sets. The most significant probe set or average probe set score may be considered for every gene.

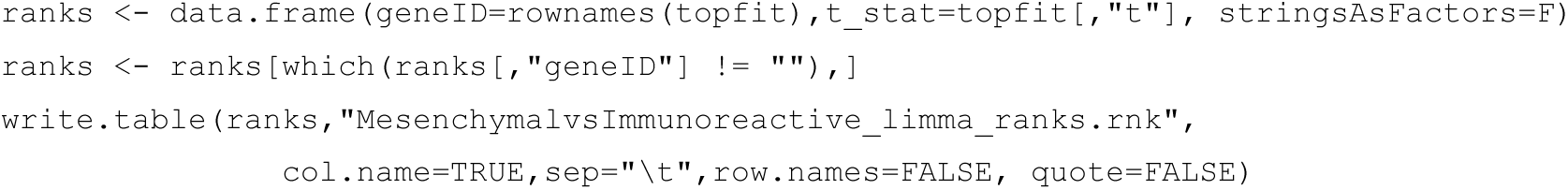

21. Create an expression file for the enrichment map and save files to the home folder of the analysis. The expression file contains the gene IDs as the first column, gene description as the second column, and the expression values for each sample as the additional columns. Gene IDs should correspond to the first column of the rank file. (Optional: biomart can be used to collect the proper names of all the genes in the expression file.) The text files will be saved on your computer in the directory specified at the beginning of the script using setwd(). The .rnk, .cls and .txt files are all tab delimited files that can be viewed in spreadsheet or in a text editor.

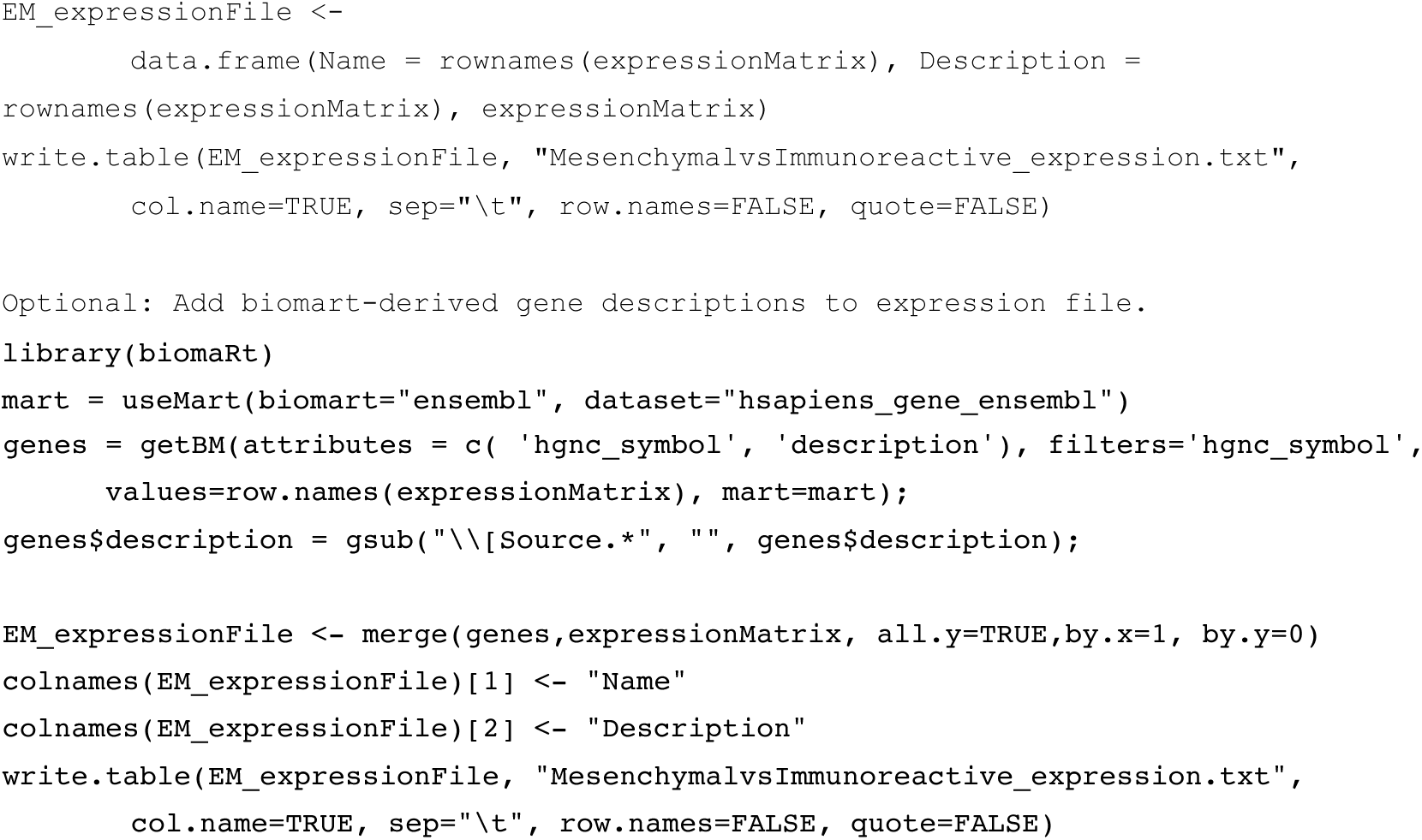

### Supplementary Protocol 3 – Pathway Enrichment Analysis in R using Roast and Camera

This protocol will demonstrate the use of R packages Roast and Camera to automate pathway enrichment analysis. Each method requires an expressionSet that minimally contains a matrix of expression values for a set of genes and conditions. The expression matrix generated in supplementary protocol part 1 or 2 is suitable for the analysis.

22. Load required Bioconductor packages into R and set working folder to the location of **Supplementary Files 1-4.**

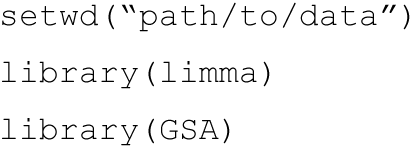

23. Load in the gene sets from a GMT file - download the latest pathway definition file automatically through R or manually through the website. Only the databases of pathway gene sets for human, mouse and rat genes are currently available on the baderlab.org downloads site. If you are working with rat or mouse data change the value of gmt_url below to specify the correct species. Consult http://download.baderlab.org/EMGenesets/current_release/ to see all available species.

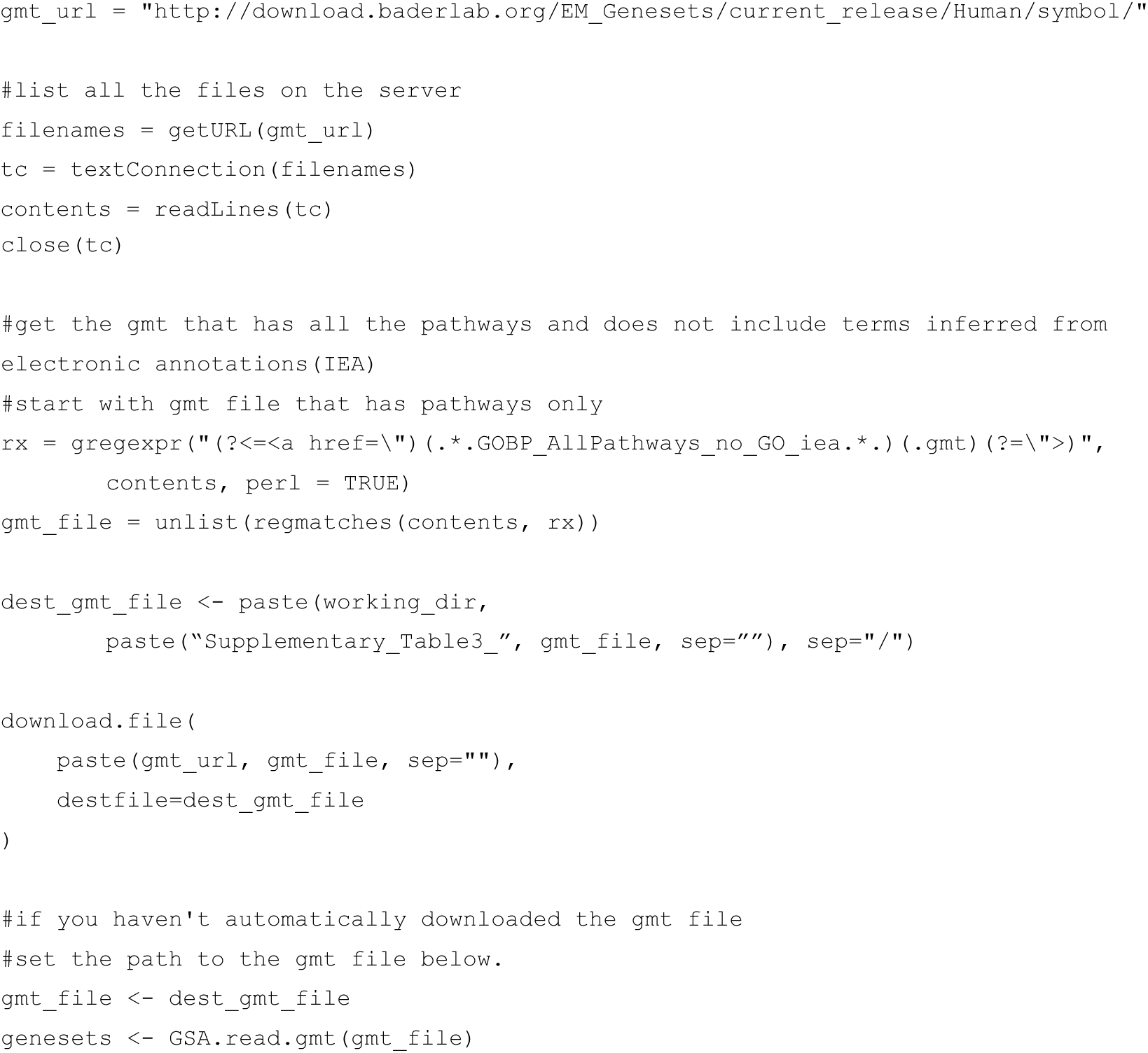

24. Camera and Roast expect the pathway gene sets to be a list of vectors where the slot name of each vector corresponds to the pathway gene set identifier, i.e. the name of the pathway, however the GSA.read.gmt() method loads the GMT file as an object with a list of pathway names and a list of pathway gene sets. Add the pathway names to the pathway gene sets vector to create a list of vectors required by Roast and Camera.

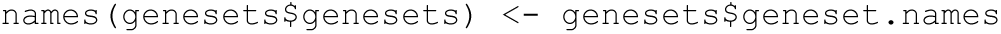

25. Specify the expression data to be used for the analysis. You can use the DGEList variable d from **Supplementary Protocol 1 (Follow steps 1-6 in Supplementary Protocol 1 to regenerate**) or MinimalSet from **Supplementary Protocol 2 (Follow steps 9-13 in Supplementary Protocol 2 to regenerate**) or. For our RNA-seq data, each row of the expression set is annotated with gene symbol and EntrezGene ID separated by “|”. To match the gene set file we need to remove the EntrezGene IDs from the row names. We choose to use gene symbols to simplify interpretation of enriched pathways and associated genes.

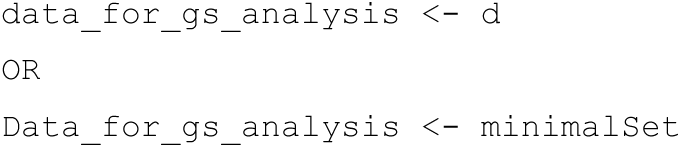

26. Camera and Roast require that the pathway gene sets are filtered such that all genes in each set have expression values in the data. Use the ids2indices() function in limma to convert gene identifiers in the gene set to indices in the data.

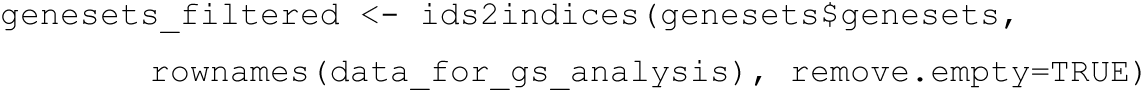

27. Filter the pathway gene sets according to their size, following the previous step of filtering by availability of expression data. Here we only include sets with more than or equal to 15 and less than 200 genes.

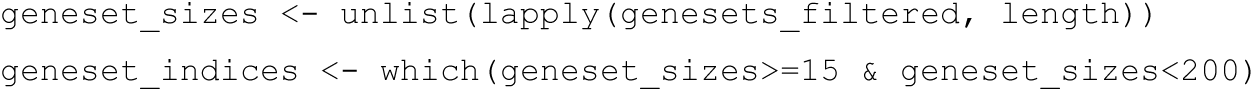

28. Create the design matrix and contrast we want to test for. In this example, we are looking for differential pathways between the Mesenchymal and Immunoreactive subtypes.

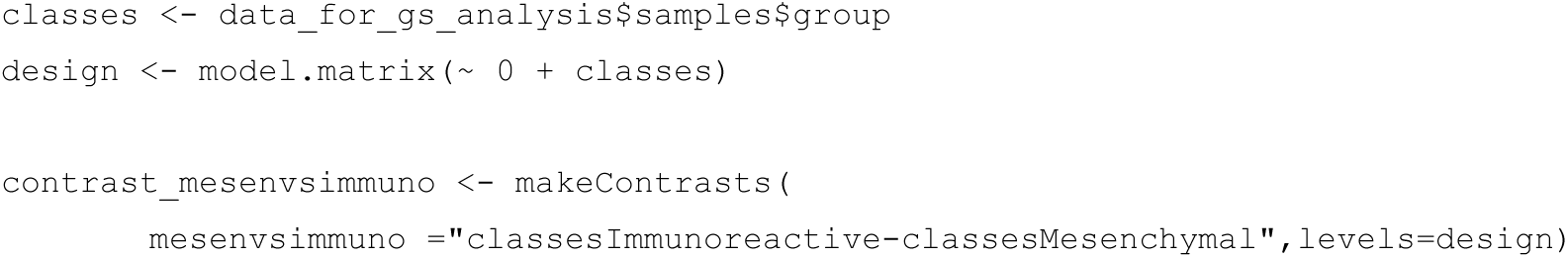

29. Run enrichment analysis and format the results to the ‘generic’ file format of Enrichment Map. This is a tab-delimited file that includes a pathway gene set name, pathway description, p-value, FDR *q*-value, phenotype and a comma-separated list of associated genes for every detected pathway. Depending on your data size and computer speed, this command could take from a few minutes to an hour to run. If you receive the warning “`In dnbinom(q, size = size, mu = mu, log = TRUE): non-integer x`”, the software has encountered unexpected non-integer values of gene expression, often indicating problems with upstream analysis such as suboptimal pre-processing or normalization procedures. Simply rounding the gene expression values may fix the error, however it should be investigated further to ensure no errors with the workflow.

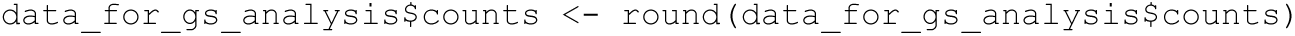

The following commands will derive results from Roast.

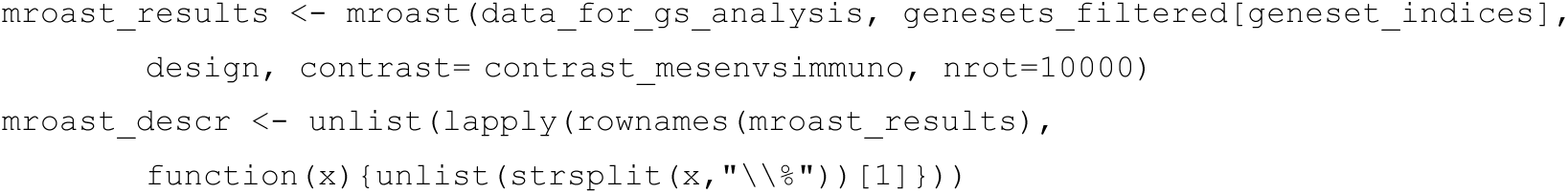

30. Inspect the results returned from Roast. The column "Direction" shows whether the gene set is enriched for up- or down-regulated genes. To ensure compatibility with Enrichment Map, convert these values such that 1 represents up-regulated and -1 represents down-regulated.

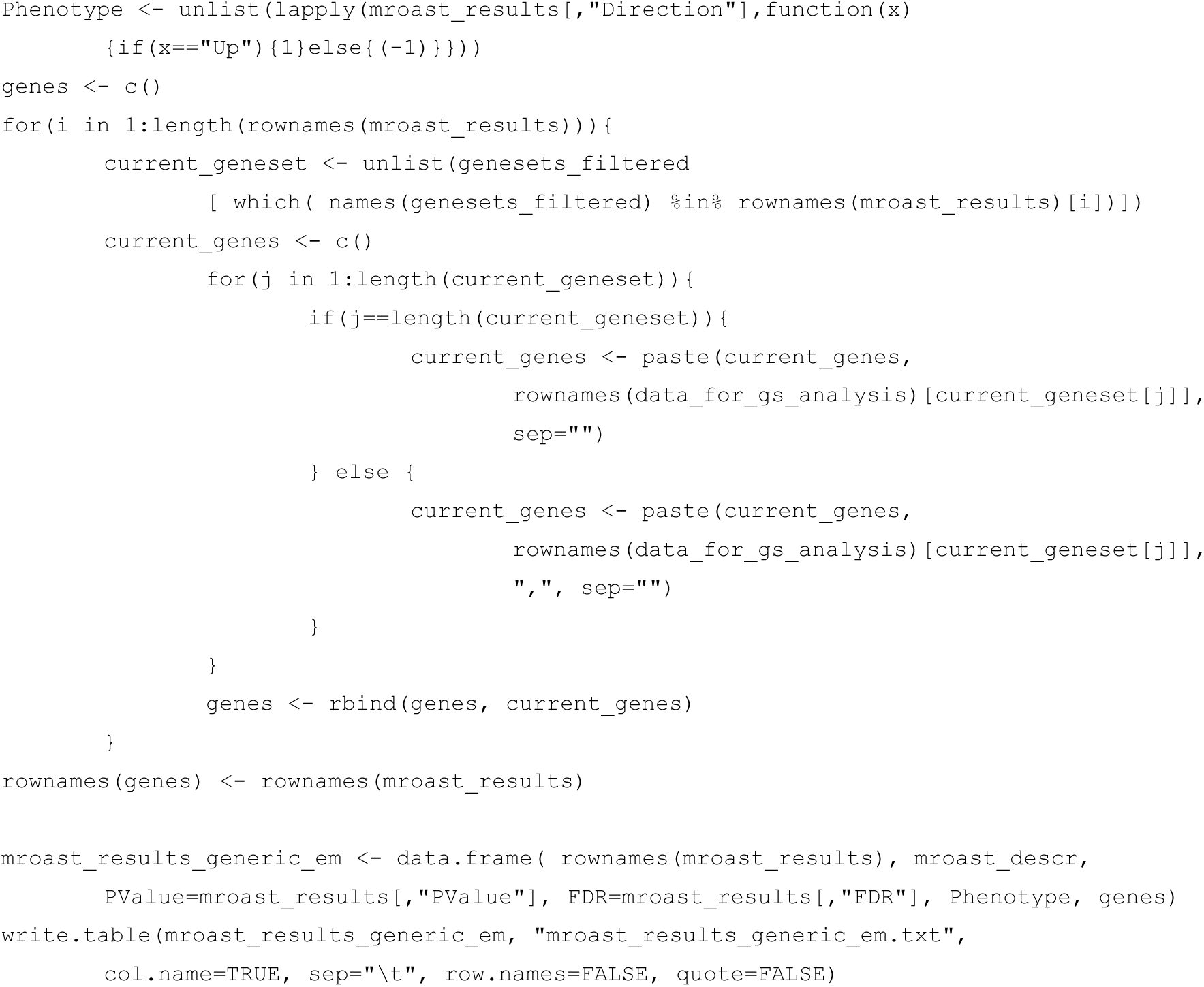

31. Run pathway enrichment analysis with the Camera R package. The analysis starts with the same files as Roast (see first four steps of **Supplementary Protocol 3**).

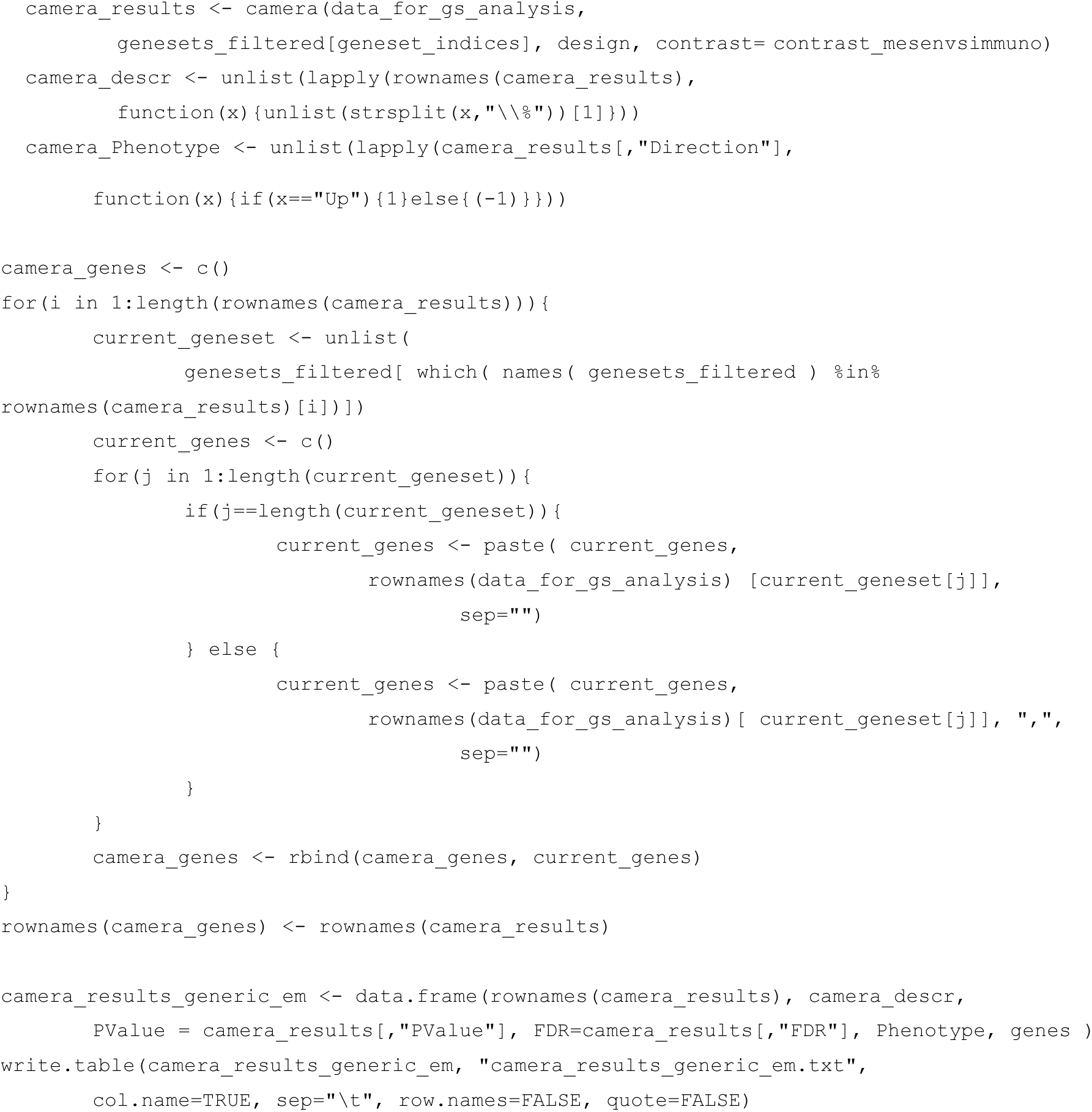

32. The results from Camera or ROAST can be input to Enrichment Map, following the protocols in the main text.

